# iPSC modeling of pulmonary arterial hypertension to uncover pathomechanisms and unrecognized modes of action of sotatercept

**DOI:** 10.64898/2026.03.12.711267

**Authors:** Aileen Schmidt, Laurien Czichon, Leonie Malhofer, Gioia Bartsch, Clara Plötner, Yuepeng Wang, Carola Voss, Anna Kuleshova, Tim Kohrn, Julia Baldauf, Astrid Weiß, Ralph Schermuly, Arjang Ruhparwar, Jan C. Kamp, Marius M. Hoeper, Ulrich Martin, Ruth Olmer

## Abstract

Pulmonary arterial hypertension (PAH) is a potentially fatal disease characterized by obliterative remodeling of distal pulmonary arteries, commonly associated with bone morphogenetic receptor type 2 (BMPR2) gene mutations. In patients with PAH, sotatercept, an activin signaling inhibitor, improves hemodynamics and outcomes, but clinical responses vary ranging from rapid improvements within weeks to no clinical benefit, suggesting additional mechanisms beyond its anti-proliferative and pro-apoptotic effects.

Using patient-specific induced pluripotent stem cell-derived smooth muscle cells (iSMCs) with BMPR2 extracellular- or kinase-domain mutations, we reproduced Activin A-driven PAH traits, including hyperproliferation, reduced apoptosis, enhanced contraction and excessive matrix production.

We identified smooth muscle cell-to-myofibroblast transition as a previously unknown contributor to pulmonary vascular remodeling and demonstrated its inhibition by sotatercept.

Beyond its established effects, sotatercept rapidly reduced contractility, collagen–integrin mechanotransduction and TGFβ receptor expression, disrupting a pathological positive feedback loop, reflected by lower levels of circulating TGFβ1 in patients on sotatercept.

Taken together, our patient-derived iSMC platform links mutation-dependent mechanisms of pulmonary vascular remodeling to variable drug responsiveness and reveals previously unrecognized, potentially rapid-acting modes of sotatercept in PAH.

**Graphical abstract:** 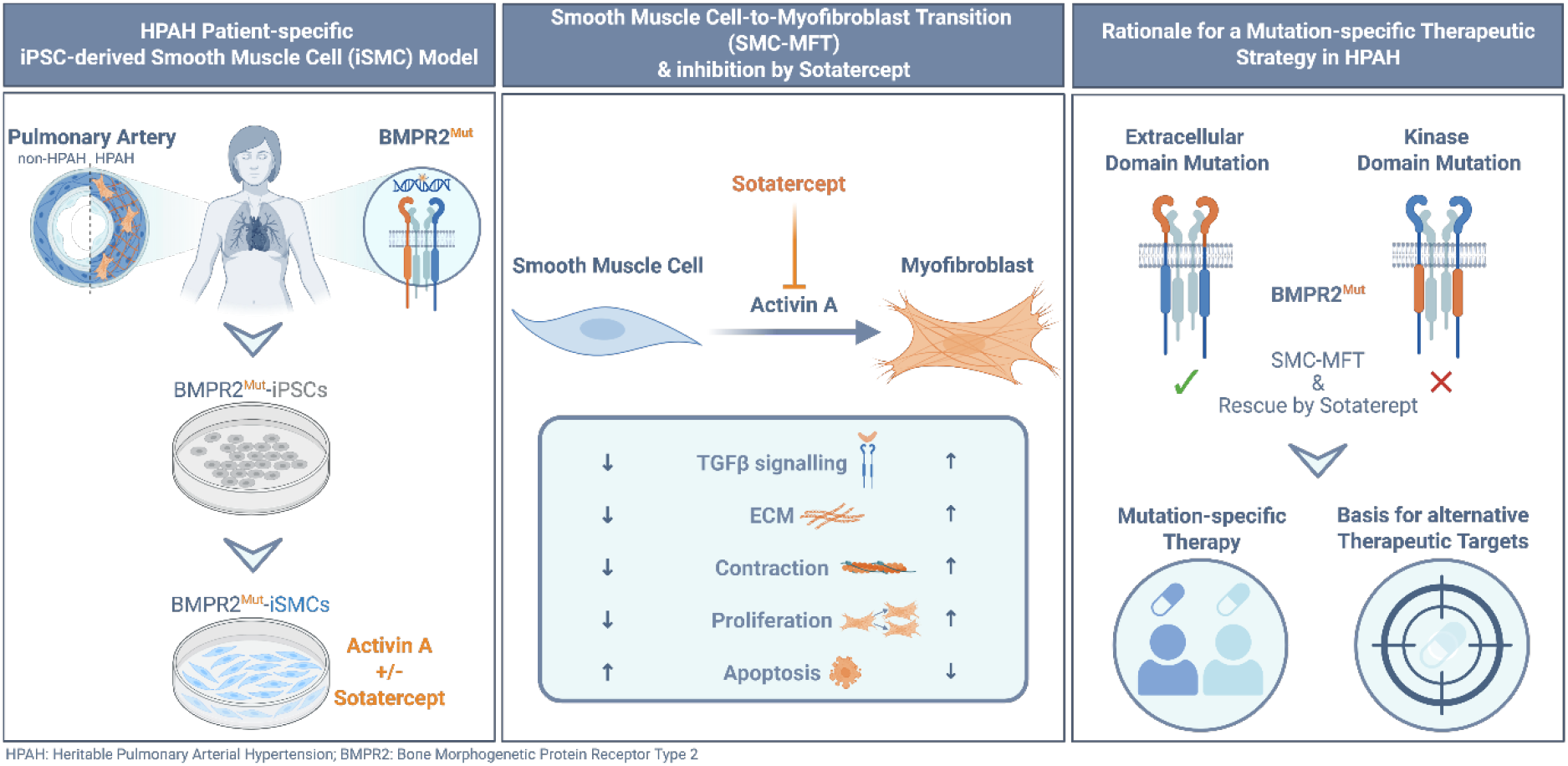

## Introduction

Heritable pulmonary arterial hypertension (HPAH) is a severe lung vasculopathy involving narrowed lumens, increased resistance and blood pressure within the pulmonary vasculature. This leads to elevated right ventricular afterload and, over time, right heart failure ^1^. HPAH is driven by pulmonary vascular remodeling, which is associated with hyperproliferation ^2^, apoptosis resistance ^3^, hypercontraction ^4^, and dysregulated extracellular matrix (ECM) deposition ^5,6^ of vascular cells.

In HPAH, pathogenic variants in the bone morphogenetic protein receptor type 2 (BMPR2) encoding gene are present in 80% of patients ^7^, leading to more severe symptomatology, increased mortality, and higher need of lung transplantation ^8^ compared to patients with idiopathic PAH. BMPR2 deficiency results in maladaptation of the Activin / TGFβ signaling, which subsequently leads to an imbalance between the canonical SMAD and SMAD-independent pathways ^9^. However, the penetrance of *BMPR2* mutations is incomplete. Only about 20–30% of carriers manifest the disease, suggesting that a single genetic defect is insufficient for disease development. The observation that only a minority of individuals with predisposing genetic mutations develop PAH has supported the second hit hypothesis in PAH, according to which second hits (e.g., inflammation, hypoxia or drug exposure) a initiate the pathological process via production of Activin A by immune cells and endothelial cells ^10^, trigger endothelial dysfunction, promote abnormal smooth muscle proliferation, and contribute to vascular remodeling, ultimately resulting in PAH ^11^. At the molecular level, Activin A and BMP 9/10, which can activate SMAD2/3 signaling via binding to a heteromer of ACVR2A/B and TGFβ type-I-receptors (Activin receptor like kinases, ALKs), are key triggers during the initiation and maintenance of the pathological processes underlying PAH ^12^.

Sotatercept, a novel drug recently approved by the FDA for the treatment of PAH, is a recombinant fusion protein functioning as an Activin/ BMP ligand trap which inhibits Activin/ BMP signaling and rebalances canonical SMAD signaling ^13^. This mitigates hyperproliferation and apoptosis resistance in vascular cells and may reverse pulmonary vascular remodeling processes ^14,15^. In clinical studies, sotatercept treatment has shown beneficial effects on haemodynamics, right heart function, and clinical outcomes, with improvement in clinical parameters and endpoints as early as week 3, reaching statistical significance by week 24 after therapy initiation ^1,16^. Although preclinical studies suggest that sotatercept’s efficacy may be driven at least in part by reverse-remodeling effects, sotatercept’s mechanism of action (MoA) in human disease is still incompletely understood. In addition, individual treatment response to sotatercept is variable ranging from normalization or near-normalization of pulmonary hemodynamics to no improvement at all. This heterogeneous response may reflect the contribution of varying pathomechanisms that depend on individual pathogenic mutations and may also suggest unrecognised MoAs of sotatercept.

Due to the limited number of patients and their heterogeneity regarding disease status, medication, co-morbidities, age, gender, nutrition, and other factors, clinical data alone will not sufficiently enable a comprehensive understanding of the pathomechanisms of PAH and the observed variations in drug responsiveness. Various *in vitro* and *in vivo* PAH models have been developed over time; however, they all possess significant limitations ^17^ and suffer from limited translational relevance ^18^.

To overcome these shortcomings, we have developed a novel *in vitro* model based on patient-derived induced pluripotent stem cells (iPSCs) that harbor mutations in various domains of the *BMPR2* gene. In contrast to recently established iPSC models of PAH that focus on endothelial cells (ECs) ^19,20^, our model builds on iPSC-derived smooth muscle cells (iSMCs), which we consider the key cell type involved in the arterial remodeling characteristic of PAH ^21^. Taking advantage of the defined conditions of this *in vitro* model, we aim to enhance our understanding of the molecular pathophysiology of PAH and to explore potential unrecognized MoAs of sotatercept. This approach intends to rationalize the heterogeneous responses to treatment observed among PAH patients and to pave the way for a more knowledge-based and individualized approach to PAH therapy.

## Results

### BMPR2^Mut^-iSMCs show PAH hallmarks of increased proliferation and decreased apoptosis, and recapitulate clinical drug responsiveness

By applying a multistep differentiation protocol, iSMCs were generated from healthy iPSCs (wildtype, WT) and from PAH patient-derived iPSCs, which carry heterozygous BMPR2 mutations. This enabled the modelling of HPAH phenotypes *in vitro* (Fig. 1 B). To exclude clonal artefacts and represent biological variability, three independent WT iPSC lines (BMPR2^WT^) derived from healthy individuals were utilized, alongside three iPSC lines generated from two patients carrying either a mutation in the extracellular domain (BMPR2^MutExDo^, heterozygous in-frame-deletion c.248-1 – 418+1 of exon 3 in the extracellular domain) or a mutation in the kinase domain (BMPR2^MutKiDo^, heterozygous missense mutation c.1471C>T in exon 11 in the kinase domain), which have recently been characterized ^22^. Highly enriched iSMCs generated from BMPR2^WT^ or BMPR2^Mut^-iPSCs demonstrated high expression levels of the SMC marker CD140b and absence of cells expressing the pluripotency associated marker OCT3/4 (Fig. 1 C; Supplementary Fig. 1 & 2). Both, BMPR2^WT^ and BMPR2^Mut^-iSMCs recapitulated typical characteristics of vascular SMCs, shown by the expression of contractile components such as alpha-smooth muscle actin (αSMA), calponin-1, and myosin heavy chain 11 (myosin-11) (Fig. 1 D, E). Expression of Myosin-11 excluded a fibroblast-like phenotype and absence of CD31 expressing cells excluded the presence of endothelial cells thereby confirming SMC identity (Supplementary Fig. 3). Global comparison of the generated iSMCs to publicly available expression data sets of primary vascular SMCs demonstrated a high degree of similarity between iSMCs and isolated primary SMCs (pSMCs), as quantified by a SMC score (Fig. 1 F). Principal Component Analysis (PCA) of RNAseq data from BMPR2^WT^ cell lines and BMPR2^Mut^-iSMCs clones revealed clustering within respective groups (Fig. 1 G). Nonetheless, the BMPR2^WT^ and BMPR2^Mut^ groups clustered separately, indicating possible HPAH-related differences in gene expression patterns.

**Figure 1:**
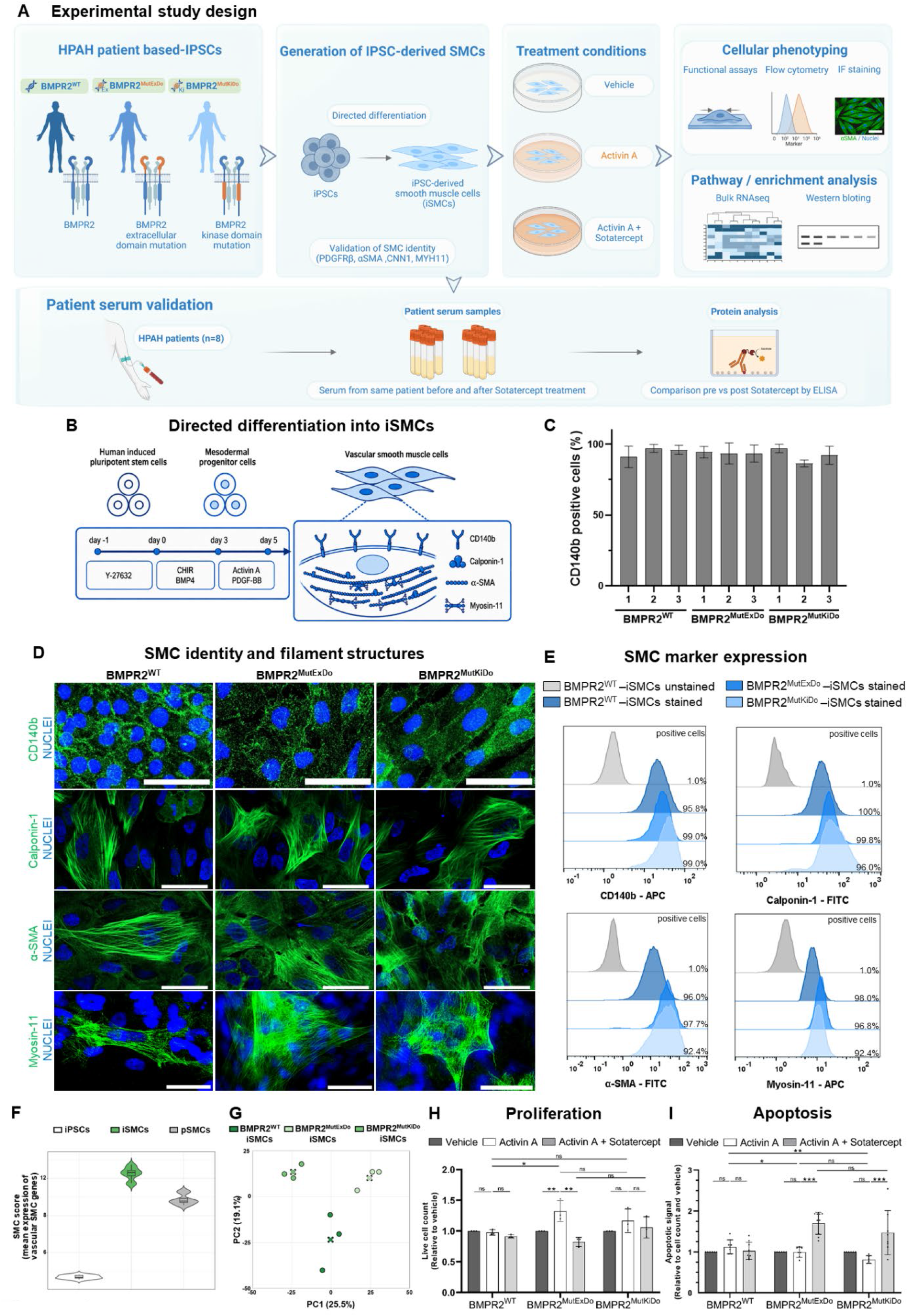
BMPR2^Mut^ iSMCs exhibit increased proliferation and decreased apoptosis, hallmarks of PAH, and recapitulate clinical drug responsiveness in vitro. (A) Schematic overview of the experimental study design. iPSC-derived SMCs carrying BMPR2^WT^, BMPR2^MutExDo^ or BMPR2^MutKiDo^ were exposed to Activin A alone or in combination with sotatercept and subsequently subjected to functional and molecular analyses. (B) Schematic illustration of chemically defined differentiation protocol from iPSCs to iSMCs. (C) Differentiation efficacy determined by flow cytometric analysis of CD140b (PDGFRβ) positive cells. (Data is presented as mean ± SD, n= 3). (D, E) Immunofluorescence staining and flow cytometry quantification confirm SMC identity of iSMCs. (D) iPSC-derived SMCs exhibit typical structures of SMC filaments as demonstrated by IF stainings with the following SMC marker panel: CD140b (PDGFRβ), alpha-smooth muscle actin (αSMA), calponin-1, myosin heavy chain 11 (myosin-11)) (green), nuclei (DAPI, blue), z-stack, scale bar 50 µm. (E) Flow cytometric characterization of iSMCs via staining for CD140b (PDGFRβ), α-smooth muscle actin (α-SMA), calponin-1 and myosin-11 (blue) relative to the unstained control (grey). Percentages indicate marker-positive cells. APC, allophycocyanin; FITC, fluorescein isothiocyanate. (F) SMC score calculated as the mean expression of Travaglini SMC genes (MSigDB: M41672) in human pulmonary artery smooth muscle cells (GSE144274), human iPSC lines (HipSci resource) and merged expression values of untreated iSMCs from all lines and clones from bulk RNA-seq. (G) PCA analysis of untreated iSMCs based on 2000 most variable genes (Log2(TPM+1)) of three independent WT cell lines (dark green dots) and three clones per BMPR2-mutant of, whereas BMPR2^MutExDo^ (light green dots) and BMPR2^MutKiDo^ (medium green dots); average of each experimental groups depicted as cross. (H) Quantification of live cell numbers relative to the respective vehicle control after 8 days of treatment. Data is presented as mean ± SD, n= 3 independent biological replicates per column indicated as dots. Exploratory two-way ANOVA with genotype and treatment as factors was followed by Sidak’s multiple-comparisons test to compare genotype differences within Activin A treatment and to compare effects of Activin A, and Activin A in combination with sotatercept, within each genotype. (I) Quantification of apoptotic signal measured as caspase3/7 activity and normalization to respective cell number and vehicle control after 5d treatment. Data is presented as mean ± SD, n= 6-9 independent biological replicates per column indicated as dots. Two-way ANOVA with genotype and treatment as factors was followed by Sidak’s multiple-comparisons test to compare genotype differences within Activin A treatment and to compare effects of Activin A and Activin A in combination with sotatercept within each genotype. ns, not significant; *p < 0.05, **p < 0.01, ***p < 0.001.

An increase in proliferation coupled with a decrease in apoptosis is recognized as hallmarks of PAH ^23^ and have previously been observed across various models ^24^. To verify whether our model recapitulates these *in vitro* phenotypes, proliferative activity and apoptosis rates following Activin A stimulation were analyzed in iSMCs derived from WT and BMPR2^Mut^ iPSC lines. Activin A, a ligand in the TGFβ signaling pathway, is elevated in PAH patients and is a known trigger of PAH disease progression ^9^. Under vehicle conditions, proliferation differed significantly between genotypes, with BMPR2^MutExDo^ iSMCs showing increased proliferation compared with BMPR2^MutKiDo^ and BMPR2^WT^ iSMCs (Supplementary Fig. 6 A). While BMPR2^MutKiDo^ iSMCs exhibited only a trend towards increased proliferation compared to WT cells, BMPR2^MutExDo^ iSMCs showed significantly elevated proliferation upon stimulation with Activin A (Fig. 1 H). Furthermore, no significant mutation-dependent differences in apoptosis resistance were detected under vehicle conditions, whereas Activin A stimulation revealed a significant reduced caspase activity in both variants of BMPR2^Mut^ iSMCs compared to BMPR2^WT^ iSMCs (Fig. 1 I; Supplementary Fig. 6 B). Based on recently collected clinical data concerning the recombinant Activin receptor 2A fusion protein, sotatercept ^16^, our objective was to further validate our *in vitro* system, particularly regarding drug responsiveness. A significant reduction in proliferation and an increase in apoptosis after co-treatment of Activin A-stimulated BMPR2^Mut^ iSMCs with sotatercept reaffirmed the established anti-proliferative and pro-apoptotic MoA of this drug, consistent with observations in other models ^25^ (Fig. 1H, I).

The forementioned results from functional assays were corroborated by gene set enrichment analyses based on bulk RNAseq data derived from generated iSMCs (Supplementary Fig. 4, 5). An inhibitory effect of sotatercept on PAH associated pro-proliferative genes, such as MYC proto-oncogene (MYC) ^26^, marker of proliferation (MKI67) ^27^, and proliferating cell nuclear antigen (PCNA) ^28,29^, in BMPR2^Mut^ iSMCs was confirmed, particularly for BMPR2^MutExDo^ iSMCs (Suppl. Fig. 4). Additionally, PAH associated pro-apoptotic genes, including caveolin-1 (CAV-1) ^30^ and caspase-4/9 (CASP4/9) ^31^, were upregulated by sotatercept in BMPR2^Mut^ iSMCs (Suppl. Fig. 5). These findings further underline that our PAH iSMC model recapitulates characteristic PAH phenotypes and provides a suitable platform for conducting drug response studies *in vitro*.

### Inhibition of Activin A-induced SMC contraction - an unrecognized MoA of Sotatercept

Hypercontractile SMCs are a central contributor to vessel stiffness and disease progression in patients with PAH^4^. By employing a functional contraction assay, the vasoconstrictive phenotype of the iSMCs was examined (Fig. 2 A, B). BMPR2^MutExDo^ iSMCs showed increased contractility under vehicle conditions compared with BMPR2^WT^ and BMPR2^MutKiDo^ iSMCs, indicating a mutation-dependent effect (Supplementary Fig. 6 C). Activin A stimulation triggered an even stronger contractile response in both BMPR2^Mut^ iSMCs than in BMPR2^WT^ iSMCs, thereby recapitulating the proposed *in vivo* phenotype of increased vasoconstriction associated with PAH^32^ (Fig. 2 A, B).

**Figure 2:**
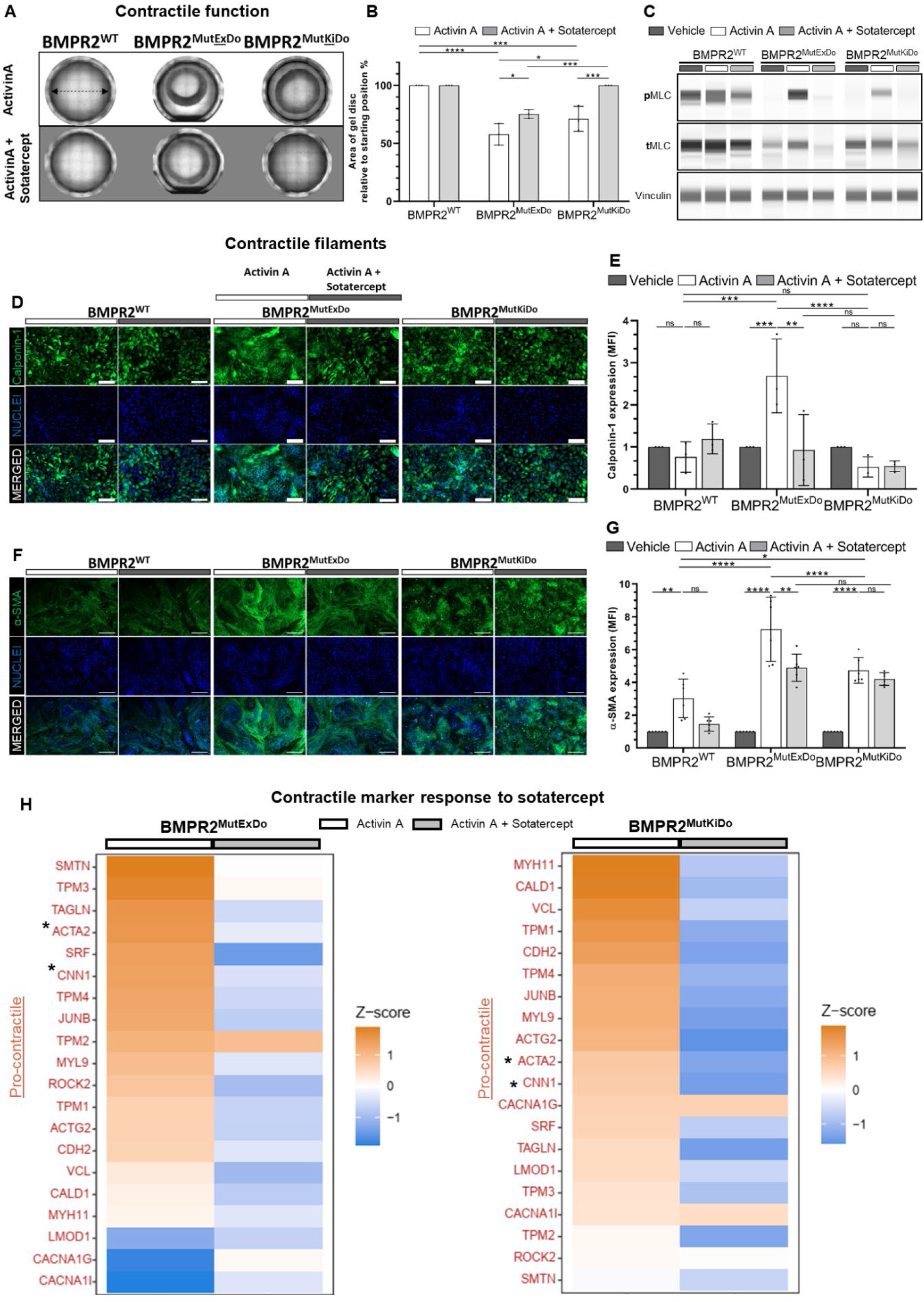
Inhibition of Activin A-induced SMC contraction by sotatercept. (**A**) Functional contraction assay with iSMCs derived from representative WT cell line or BMPR2^Mut^ clones. Double arrow indicates well diameter as starting position for contraction. (**B**) Contraction was determined as the area of the gel disc after 72 h relative to the initial area of the iSMC-gel-discs. Data is presented as mean ± SD, n= 3 independent biological replicates per column indicated as dots. Exploratory two-way ANOVA with genotype and treatment as factors was followed by Sidak’s multiple-comparisons test to compare genotype differences within Activin A treatment and to compare effects of Activin A and Activin A in combination with sotatercept within each genotype. ns, not significant; *p < 0.05, **p < 0.01, ***p < 0.001. (**C**) Exemplary analysis of phosphorylated myosin light chain (pMLC) and total MLC as a readout of actomyosin activation and contractile signaling by automated capillary-based western blot analysis for each genotype and treatment condition (vehicle, Activin A, Activin A + Sotatercept). (**D, F**) Immunofluorescence staining of components of the contractile apparatus in iSMCs of one iPSC line or clone representative for the respective genotype (BMPR2^WT^, BMPR2^MutExDo^, BMPR2^MutKiDo^). Calponin-1 and αSMA (green), nuclei DAPI (blue), scale bar 200 µm. (**E**) Quantification of calponin-1 from flow cytometry analysis (MFI) relative to the respective vehicle control. Data is presented as mean ± SD, n= 3 independent biological replicates per column indicated as dots. Exploratory two-way ANOVA with genotype and treatment as factors was followed by Sidak’s multiple-comparisons test to compare genotype differences within Activin A treatment and to compare effects of Activin A and Activin A in combination with sotatercept within each genotype. ns, not significant; *p < 0.05, **p < 0.01, ***p < 0.001. (**G**) Quantification of αSMA from immunofluorescence images expressed as mean fluorescence intensity (MFI) normalized to cell number (DAPI) and relative to the respective vehicle control. Data is presented as mean ± SD, n= 6 independent biological replicates per column indicated as dots. Two-way ANOVA with genotype and treatment as factors was followed by Sidak’s multiple-comparisons test to compare genotype differences within Activin A treatment and to compare effects of Activin A and Activin A in combination with sotatercept within each genotype. ns, not significant; *p < 0.05, **p < 0.01, ***p < 0.001. (**H**) Heatmaps showing differentially expressed genes with pro-contractile function in Activin A vs. Activin A + sotatercept treated iSMCs each condition normalized to the untreated control. For contraction gene set scoring (MSigDB: M1429), bulk RNA-seq expression values were analysed using a gene-by-sample FPKM matrix (mean FPKM of three independent biological replicates per group, Activin A and Activin A + sotatercept). Stars (*) highlight mentioned genes in the respective text.

The contractile activity of SMCs is directly influenced by the level of phosphorylated myosin light chain (pMLC), which activates the actin-myosin cross-bridge cycle, and is supposed to be alleviated in PAH patients ^33,34^. Western blot analysis of phosphorylated and total MLC indicated an increased proportion of pMLC in Activin A stimulated BMPR2^Mut^ iSMCs compared to Activin A-stimulated BMPR2^WT^ iSMCs, (Fig. 2 C). Interestingly, the Activin ligand trap sotatercept has not been associated with any effects on vasoconstriction *in vitro* studies, animal studies, or clinical trials. Consequently, we investigated such effects within our HPAH iSMC model. Remarkably, western blotting suggested that co-administration of sotatercept may also reduce proportion of pMLC in Activin A stimulated BMPR2^Mut^ iSMCs (Fig. 2 C).

To further elucidate the effect of sotatercept on SMC contractility, the composition of the contractile apparatus was analyzed, including actin filaments (αSMA) and actin-bound calponin-1, which are essential for the fine-tuning of actin-myosin cross-bridging ^35^. Immunofluorescence (IF) staining and protein quantifications demonstrated elevated calponin-1 in BMPR2^MutExDo^ iSMCs after Activin A treatment compared to WT cells, which was not observed in BMPR2^MutKiDo^ iSMCs (Fig. 2 D, E). Consistent with the contraction assay results, αSMA was upregulated in BMPR2^Mut^ iSMCs following Activin A treatment, with a particularly pronounced increase in BMPR2^MutExDo^ iSMCs (Fig. 2 F, G). Furthermore, in line with the contraction assay findings, sotatercept significantly reduced levels of calponin-1 and αSMA in Activin A-stimulated BMPR2^MutExDo^ iSMCs. However, even with sotatercept treatment αSMA levels in BMPR2^MutExDo^ iSMCs remained the highest across the genotypes (Fig. 2 D – G). In BMPR2^MutKiDo^ iSMCs, sotatercept had no effect on the already low calponin levels in Activin A-stimulated cells and induced only a modest downward trend in αSMA expression, which remained above WT levels.

A gene set enrichment analysis was performed to further explore the molecular mechanisms underlying the effect of sotatercept on SMC contraction (Fig. 2 H). Although the effect was more prominent in Activin A-treated BMPR2^MutExDo^ iSMCs, sotatercept appears to exert its anti-contractile effect by downregulating the expression of positive regulators of SMC contraction. For instance, sotatercept reduced the expression of genes associated with PAH, such as myosin heavy chain 11 (MYH11) ^36^, αSMA filaments (ACTA2) ^9^, calponin-1 (CNN1) ^28^, transgelin (TAGLN) ^37^, as well as other critical genes involved in SMC contraction, including myosin light chain 9 (MYL9) ^38^ and smoothelin (SMTN) ^39^, in both BMPR2^Mut^ iSMCs.

### SMC-to-Myofibroblast transition (SMC-MFT) contributes to arterial tissue remodeling in PAH

Increased hydrodynamic parameters and vessel stiffness attributed to heightened ECM content have been documented in patients with PAH ^5^. Myofibroblasts are regarded as the main producers of ECM during the pathological tissue remodeling in PAH ^40,41^. While it has been proposed that, in the case of PAH, such cells originate from local endothelial cells via endothelial-to-mesenchymal transition (EndoMT) ^42^, or are derived from quiescent resident fibroblasts or bone marrow-derived circulating cells ^40,41^, we questioned whether myofibroblasts could also arise from our highly enriched BMPR2^Mut^ iSMCs cultures, serving as an *in vitro* counterpart of BMPR2^Mut^ local arterial SMCs. Given the significant involvement of collagen I in pathological tissue remodeling in PAH and considering that it is typically produced at elevated levels by myofibroblasts, we examined collagen I production following Activin A stimulation of BMPR2^Mut^ iSMCs. In comparison to Activin A-stimulated BMPR2^WT^ iSMCs, an increased expression of collagen I was observable in Activin A stimulated BMPR2^Mut^ iSMCs, which likewise exhibited co-expression of αSMA, a marker known to be expressed in SMCs and myofibroblasts ^43^. In particular, in Activin A-treated BMPR2^MutExDo^ iSMCs, a dense network of collagen I fibers was evident (Fig. 3 A). Quantitative analysis of collagen deposition by a SiriusRed assay revealed a significantly elevated amount of collagen I per cell in Activin A-stimulated BMPR2^Mut^-iSMCs relative to the Activin A-stimulated BMPR2^WT^iSMCs, particularly pronounced in BMPR2^MutExDo^ iSMCs (Fig. 3 B). RNASeq analyses further confirmed a substantial upregulation of various additional genes involved in ECM production, particularly in Activin A-treated BMPR2^MutExDo^ iSMCs (Fig. 3C). These included collagen I ^44^, ITGA2 ^45^, TGFβ1 ^46^, TGFβR1 ^47^, as well as ECM-associated genes such as fibronectin (FN1) ^48^ and tenascin (TNC) ^49^, all of which are linked to PAH.

**Figure 3:**
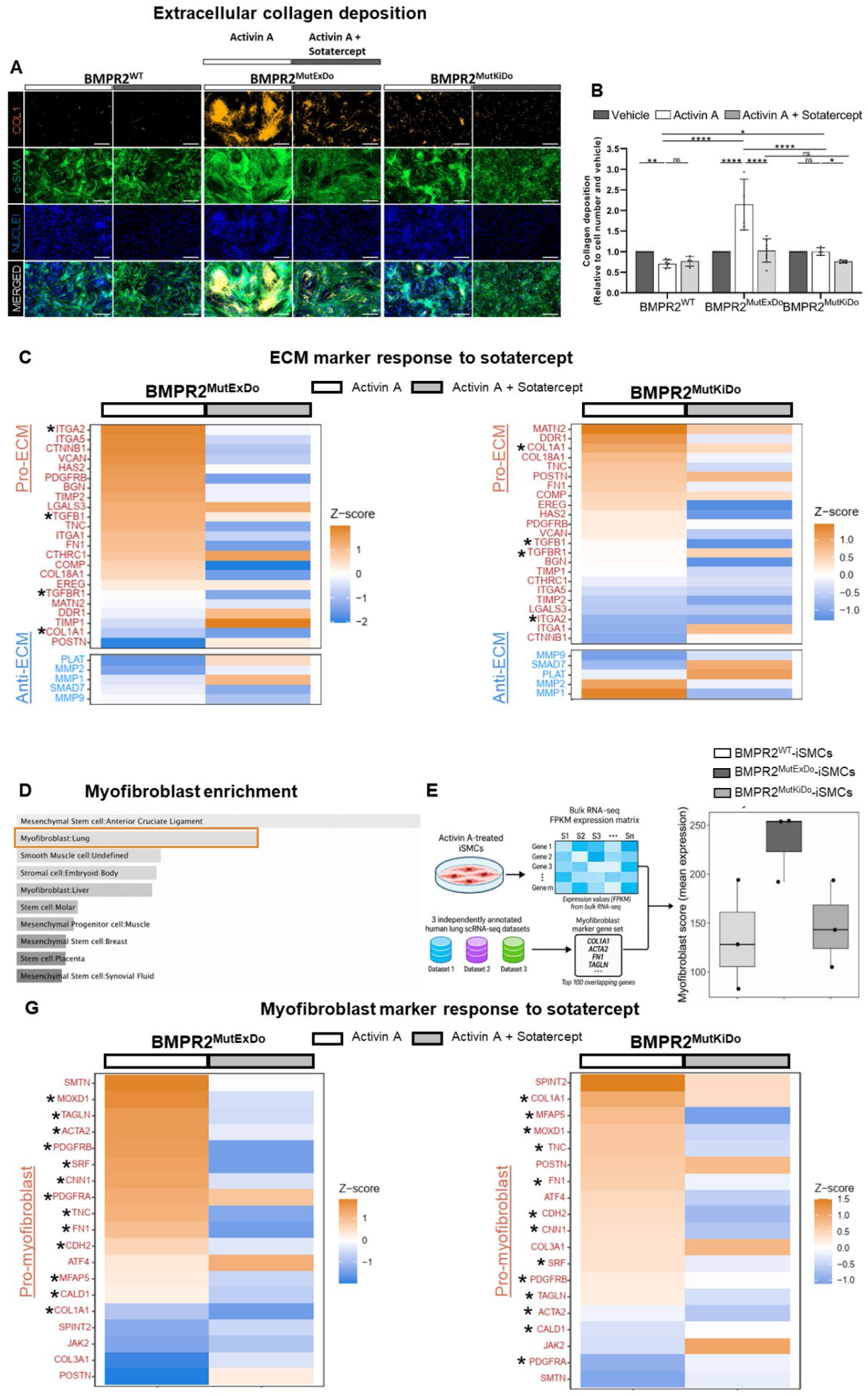
SMC-to-Myofibroblast transition (SMC-MFT) contributes to arterial tissue remodeling in PAH. (**A**) Immunofluorescence staining of iSMCs from one iPSC line or clone representative for the respective genotype (BMPR2^WT^, BMPR2^MutExDo^, BMPR2^MutKiDo^) for collagen I (orange) and αSMA (green), nuclei DAPI (blue), scale bar 200 µm. (**B**) Quantification of collagen I deposition per cell relative to the respective vehicle control by siriusRed assay. Data is presented as mean ± SD, n= 4-7 independent biological replicates per column indicated as dots. Two-way ANOVA with genotype and treatment as factors was followed by Sidak’s multiple-comparisons test to compare genotype differences within Activin A treatment and to compare effects of Activin A and Activin A in combination with sotatercept within each genotype. ns, not significant; *p < 0.05, **p < 0.01, ***p < 0.001. (**C**) Heatmaps showing differentially expressed genes associated with extracellular matrix production in Activin A vs. Activin A + sotatercept treated iSMCs each condition normalized to the untreated control. For ECM gene set scoring (MSigDB: M11366), bulk RNAseq expression values were analyzed using a gene-by-sample FPKM matrix (mean FPKM of three independent biological replicates per group, Activin A and Activin A + sotatercept). Stars (*) highlight mentioned genes in the respective text. (**D**) Functional enrichment analysis of positive regulated genes from the gene set enrichment analysis from BMPR2^Mut^ iSMCs bulk RNAseq with Enrichr ^97^. Bar plot shows enriched cell types from CellMarker 2024. Bar length represents significance of enrichment. (**E**) Schematic overview of the myofibroblast signature score analysis. Myofibroblast-associated marker genes were defined as the top 100 overlapping myofibroblast-associated marker genes from three independently annotated human lung scRNA-seq datasets: GSE135893, GSE136831 and the Munich cohort dataset from Mayr et al, 2021 ^98^. Their expression values were extracted from the bulk RNA-seq FPKM matrix of Activin A-treated iSMCs and used to calculate the myofibroblast signature score. Myofibroblast signature score from the bulk RNA-seq FPKM expression matrix of Activin A treated iSMCs. Boxes indicate the interquartile range (25th–75th percentile), the center line denotes the median, and whiskers extend to 1.5 × the interquartile range. Each point represents an individual biological sample. (**G**) Heatmaps showing differentially expressed pro-myofibroblast genes in Activin A vs. Activin A + sotatercept treated iSMCs each condition normalized to the untreated control. For myofibroblast gene set scoring, bulk RNAseq expression values were analysed using a gene-by-sample FPKM matrix (mean FPKM of three independent biological replicates per group, Activin A and Activin A + sotatercept). Stars (*) highlight mentioned genes in the respective text.

The concurrent presence of contractile and ECM features in Activin A treated BMPR2^Mut^ iSMCs prompted us to further explore the molecular signature of Activin A treated BMPR2^Mut^ iSMCs. We observed global similarities to myofibroblasts and fibroblasts in Activin A-treated BMPR2^Mut^ iSMCs from a functional enrichment analysis of bulk RNAseq data (Fig. 3 D). Additionally, an integration of three independent databases, comprising a shared set of approximately 100 genes, identified a robust enrichment of a myofibroblast signature in Activin A-treated BMPR2^MutExDo^ iSMCs. The enrichment was less pronounced in BMPR2^MutKiDo^ iSMCs when compared to Activin-treated BMPR2^WT^ iSMCs (Fig. 3 E).

Interestingly, sotatercept also rescued the elevated expression of various core myofibroblast markers, including platelet-derived growth factor receptor alpha/beta (PDGFRA/B) ^50^, αSMA filaments (ACTA2) ^41,50^, collagen type I alpha 1 chain (COL1A1) ^51^, fibronectin 1 (FN1) ^51,52^, caldesmon 1 (CALD1) ^52^, cadherin 2 (CDH2) ^52^, calponin 1 (CNN1) ^53,54^, microfibril-associated glycoprotein 5 (MFAP5) ^55,56^, tenascin C (TNC) ^54^, transgelin (TAGLN) ^57^, and monooxygenase, DBH-like (MOXD1) ^58^ in Activin A–treated BMPR2^Mut^ iSMCs (Fig. 3 G).

### Activin A induces mechano-signaling / TGFβ signaling in BMPR2^Mut^-iSMCs

It is well established that accumulated ECM proteins mechanotransduce signals via integrin complexes (ITGA/B), hereby contributing to a positive feedback loop ^59^ that promotes vascular remodeling in pulmonary hypertension ^60,61^. Therefore, we analyzed the expression of the collagen I corresponding integrin receptor subunit 2A (ITGA2) in Activin A treated BMPR2^Mut^ iSMCs. Indeed, integrin α2 expression was significantly increased in Activin A stimulated BMPR2^MutExDo^ iSMCs compared to the Activin A-stimulated BMPR2^WT^ iSMCs, while no upregulation was detectable on BMPR2^MuKiDo^ iSMCs (Fig. 4 A, B). Given the established interplay between collagen–integrin-dependent mechanotransduction, actomyosin contractility, and TGF-β signaling, we investigated modulation of TGF-β pathway activity. Interestingly, TGFβR1, which – following tetramerisation with TGFβR2 – is responsible for signal transduction via SMAD and non-SMAD signaling ^62^, was significantly upregulated in Activin A-stimulated BMPR2^MutExDo^ iSMCs compared to the Activin A stimulated BMPR2^WT^ iSMCs, similar to recent results of concurrent elevation of Activin A and TGFBR1 (ALK5) in PAH SMCs and ECs ^12^. To date, no direct evidence demonstrates that Activin A directly induces TGFBR1 (ALK5) expression in PAH-SMCs; but an indirect induction via TGF-β1^63^ generated within a feed-forward signaling loop could lead to Activin A induced TGFBR1 upregulation. Controversially, no upregulation was detectable in BMPR2^MutKiDo^ iSMCs upon Activin A stimulation (Fig. 4 C, D). Remarkably, sotatercept not only largely rescued the increased collagen I protein expression in BMPR2^MutExDo^ iSMCs but also resulted in a significant downregulation of the integrin receptor subunit 2A and TGFβR1 in BMPR2^MuExDo^ iSMCs compared to cells treated with Activin A. Such effects were not observed in BMPR2^MuKiDo^ iSMCs (Fig. 4 A – D). Given the observed elevated expression of TGFβ1 (Fig. 3 C) in myofibroblasts derived from Activin A treated BMPR2^Mut^ iSMCs, we reasoned an inhibitory effect of sotatercept. To explore whether TGFβ1 contributes to PAH severity and drug responsiveness in patients with *BMPR2* mutations, we analyzed serum TGFβ1 levels in such PAH patients before and on sotatercept therapy. Interestingly, elevated TGFβ1 levels in PAH patients significantly decreased with sotatercept therapy (Figure 4 E). The existence of the observed phenotypic features in Activin A–stimulated BMPR2^Mut^ iSMCs may be explained by the well-established and consistently recapitulated shift toward enhanced SMAD2/3 signaling ^9^, which is detectable in both mutants (Fig. 4 F and supplementary Fig. 7 A, B). Moreover, sotatercept reduced phosphorylation of SMAD2/3, consistent with previous observations in primary PAH-SMCs, while SMAD1/5/9 signalling remained largely unaffected^9^. Consequently, sotatercept may also counteract the Activin A induced and TGFβ-driven and mechano-signaling-mediated positive feedback loop involved in PAH pathogenesis (Fig. 5).

**Figure 4:**
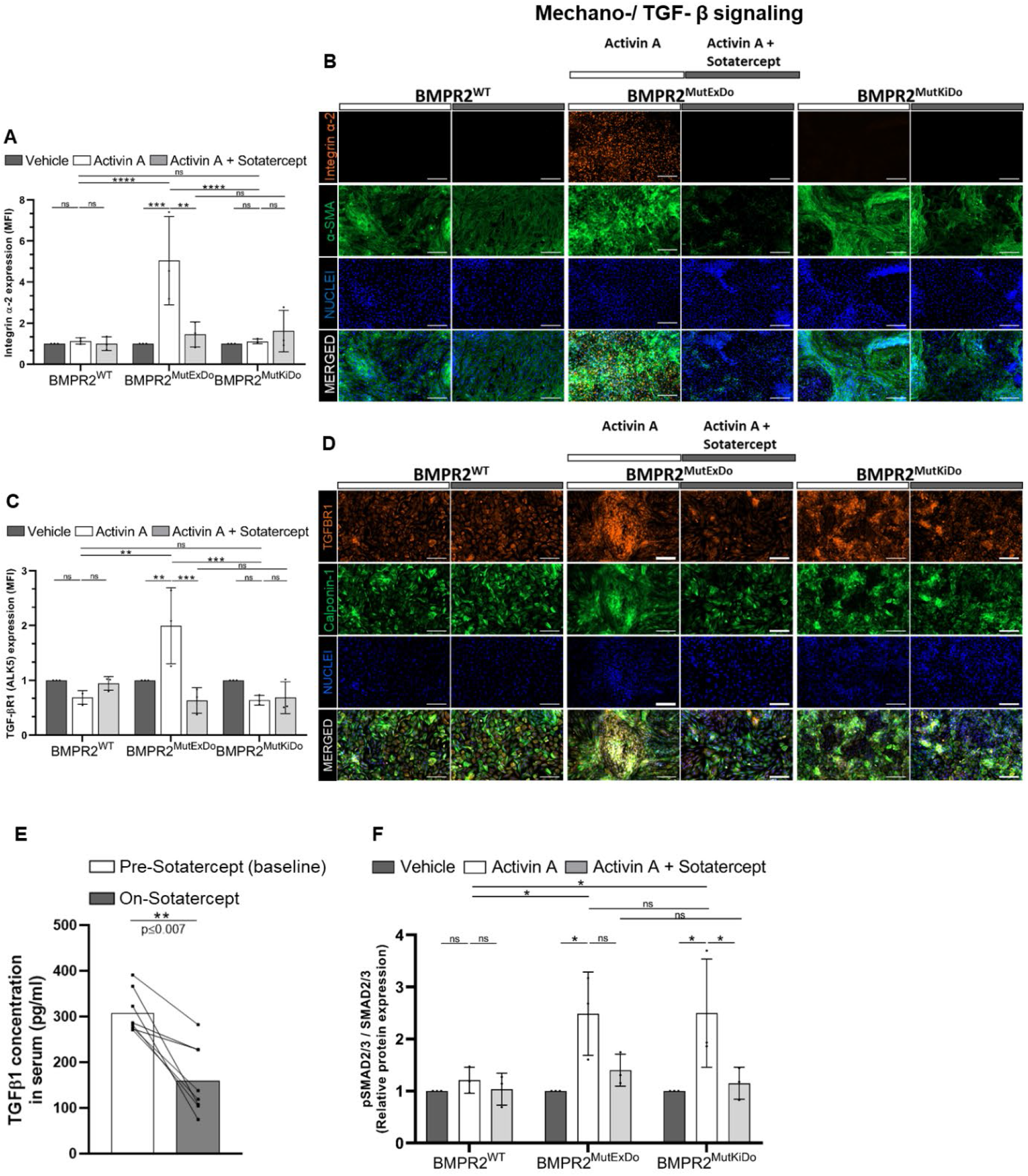
Activin A induces a pathological upregulation of collagen-integrin mechano-signaling and TGFβ-signaling in BMPR2^MutExDo^ iSMCs. (**A**) Quantification of integrin α2 iSMCs via flow cytometry (MFI). Data is presented as mean ± SD, n= 3 independent biological replicates per column indicated as dots. Exploratory two-way ANOVA with genotype and treatment as factors was followed by Sidak’s multiple-comparisons test to compare genotype differences within Activin A treatment and to compare effects of Activin A and Activin A in combination with sotatercept within each genotype. ns, not significant; *p < 0.05, **p < 0.01, ***p < 0.001. (**B**) Immunofluorescence staining of iSMCs from one line/clone representative for the respective genotype (BMPR2^WT^, BMPR2^MutExDo^, BMPR2^MutKiDo^) stained for integrin-α2 (orange), actin (green), nuclei DAPI (blue), scale bar 200 µm. (**C**) Quantification of TGFβR1 in iSMCs via flow cytometry (MFI). Data is presented as mean ± SD, n= 3 independent biological replicates per column indicated as dots. Exploratory two-way ANOVA with genotype and treatment as factors was followed by Sidak’s multiple-comparisons test to compare genotype differences within Activin A treatment and to compare effects of Activin A and Activin A in combination with sotatercept within each genotype. ns, not significant; *p < 0.05, **p < 0.01, ***p < 0.001. (**D**) Immunofluorescence staining of iSMCs derived from one iPSC line / clone representative for the respective genotype (BMPR2^WT^, BMPR2^MutExDo^, BMPR2^MutKiDo^) for TGFβR1 (orange) and calponin (CCN1, green) nuclei DAPI (blue), scale bar 200 µm. (**E**) TGFβ content in patient sera at baseline and on sotatercept treatment determined by ELISA (on-treatment samples obtained 9–41 months after sotatercept initiation). Each dot represents an individual patient (n=8), with lines connecting paired pre-and on-sotatercept. Horizontal lines indicate mean. Statistical significance was assessed using a two-tailed Wilcoxon matched-pairs sign-rank test. ns, not significant; *p < 0.05, **p < 0.01, ***p < 0.001. (**F**) Quantification of phosphorylated SMAD2/3 (p-SMAD2/3) relative to total SMAD2/3 (t-SMAD2/3), normalized to vinculin and the respective vehicle control by automated capillary-based western blot analysis. Data is presented as mean ± SD, n= 3 independent biological replicates per column indicated as dots. Exploratory two-way ANOVA with genotype and treatment as factors was followed by Sidak’s multiple-comparisons test to compare genotype differences within Activin A treatment and to compare effects of Activin A and Activin A in combination with sotatercept within each genotype. ns, not significant; *p < 0.05, **p < 0.01, ***p < 0.001.

**Figure 5:**
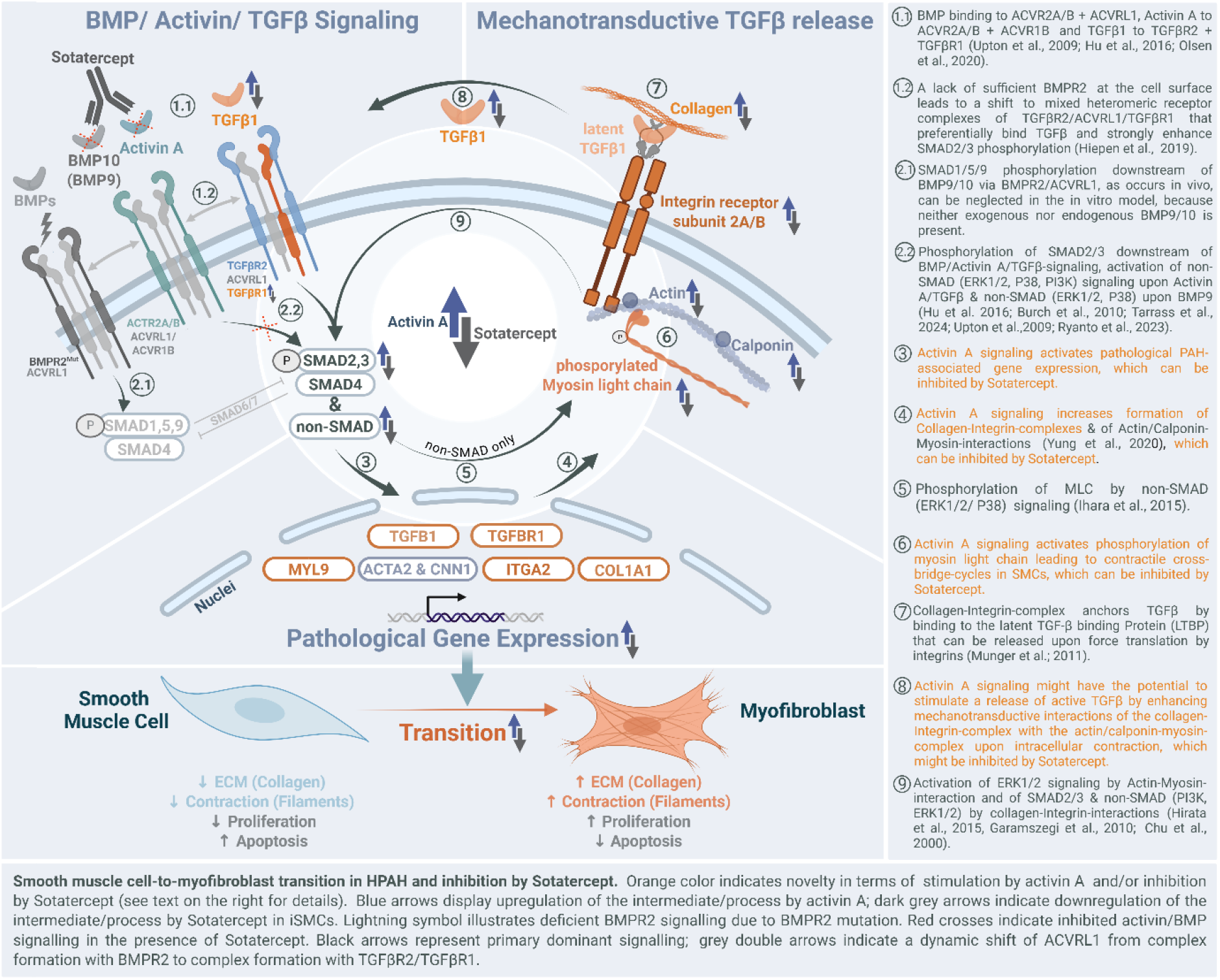
Schematic illustration of the hypothesized maladaptive Activin A/TGFβ signaling axis in crosstalk with the mechanosignaling of collagen I-integrin/actomyosin interactions leading to a pathological feedback loop, triggering smooth muscle cell–to– myofibroblast transition (SMC-MFT) in BMPR2^Mut^-iSMCs. Created in BioRender. Olmer, R. (2026) https://BioRender.com/d3lee4y

## Discussion

Despite extensive clinical and experimental research, the molecular pathomechanisms of PAH have largely remained unclear until now. Several factors have prevented a deeper understanding of the disease. It is extremely challenging to draw conclusions from clinical data, even in multicentre clinical trials, due to the relatively small and highly heterogeneous patient population, which carries different mutations and shows diverse phenotypes and drug responsiveness. PAH research is further complicated by the fact that patient’s cells and histological specimens are almost exclusively derived from few autopsies or explants after lung transplantation.

Until very recently ^64^, traditional animal models in rats, and especially mice did not reflect the severity and the genetic causes of the human disease or sex-related differences in prevalence. Even in the era of targeted genetic engineering via CRISPR-Cas, the generation of genetically engineered rats remains laborious, time-consuming, expensive, and the introduction of a multitude of numerous diverse mutations is impractical. Moreover, it is generally difficult to replicate the slow progression of PAH *in vitro*, as only minor differences between wild-type and mutated cells are to be expected during the typically short cell culture periods.

The elegance of our iPSC-based system lies in its capacity to eliminate the influence of disease stage, comorbidities, medications, environmental exposures, infections, and other confounding factors. These highly individual factors significantly complicate the analysis of clinical data, obscuring the underlying, mutation-dependent differences in dysregulated pathways and disease phenotypes. Despite the reduced complexity, our iPSC model still incorporates patient-specific genetic features, allowing us to directly correlate our experimental results with data from clinical studies. This includes the observed drug responsiveness of patients with specific mutations. The unlimited expansion potential of iPSCs poses another advantage for PAH research: Unlike primary cells, iPSCs enable the conduct of numerous repetitive experiments necessary to attain statistical significance, even when discerning minor phenotypic differences between WT and mutated cells, which are anticipated in a disease that develops gradually over years rather than days.

Based on preliminary data from our recent clinical trials, we established iPSC lines from two patients that are heterozygous carriers of pathogenic mutations either in the extracellular or in the kinase domain of the *BMPR2*, employing the same reprogramming protocol.

Despite the relatively small differences observed in functional measurements and expression levels, our experiments conducted under well-defined conditions demonstrated high reproducibility among individual cell clones derived from a single PAH patient. We intentionally minimized complexity by incorporating only one cell type – SMCs in our model. While other models have previously employed patient-specific iPSC-derived ECs ^19,20^, and addressed the role of ECs during disease initiation, we hypothesized that SMCs are central to understanding the pathological pulmonary arterial remodeling in PAH. In this study, experiments were conducted with highly pure mesoderm-derived iPSC-SMCs, the expression of Myosin-11 excluded a fibroblast-like phenotype and absence of CD31^pos^ cells excluded an EC phenotype as well as contaminating ECs confirming the presence of highly enriched SMCs. Remarkably, the SMC score of our iPSC-derived iSMCs was even higher than that of primary SMCs isolated from human pulmonary arteries, which may be attributed to contamination of primary cells with other vascular cell types, particularly ECs or fibroblasts. As expected, iSMCs differentiated from the three clones established from each patient clustered closely together, whereas the three WT lines from different donors clustered less closely.

The first pivotal step was to recapitulate the hallmarks of PAH, i.e., hyperproliferation and increased resistance to apoptosis. We confirmed increased proliferation following Activin A stimulation in iSMCs differentiated from all three BMPR2^MutExDo^ clones, compared to BMPR2^WT^ lines. Apoptosis was significantly reduced in iSMCs carrying both mutations. Interestingly, co-treatment with Activin A and sotatercept rescued both proliferation and apoptosis, approximately to WT levels.

RNAseq analyses corroborated the findings of the functional assays, as evidenced by the significant upregulation of pro-proliferative genes in BMPR2^MutExDo^ iSMCs. In contrast, BMPR2^MutKiDo^ iSMCs exhibited a notable downregulation of anti-proliferative genes. For both mutations, reduced apoptosis appears to correlate more strongly with the downregulation of pro-apoptotic genes than with the upregulation of anti-apoptotic genes. Our transcriptome analyses provide additional support for the findings related to the clinical effects of sotatercept. The concurrent administration of sotatercept seems to correct the Activin A induced dysregulation of various genes associated with proliferation and apoptosis in both mutations. The observed discrepancies between the two mutations concerning Activin stimulation and the effects of sotatercept on dysregulated genes related to proliferation and apoptosis may suggests different underlying pathomechanims and as yet unrecognized involvement of regulatory pathways beyond the canonical SMAD signaling.

Although the current study results do not definitively indicate whether sotatercept treatment can effectively reverse an already established Activin A-induced activation state of SMCs, the data nonetheless clearly recapitulate the known hallmarks of PAH and the therapeutic effects of sotatercept.

Next to hyperproliferation and resistance to apoptosis observed in vascular cells, increased vasoconstriction has been reported in PAH ^65,66^. Pulmonary vascular dysfunction is present from the early stages of pathogenesis in pre-capillary arterioles, contributing to the end stage disease characterized by obliteration of the precapillary pulmonary arterioles ^65^. The results of the conducted contraction assay also reflect the clinical PAH phenotype of increased vasoconstriction. iSMCs harboring both mutations exhibited a significantly increased contraction response compared to WT iSMCs upon Activin A stimulation. The observed elevated ratio of phosphorylated to non-phosphorylated myosin light chain (MLC) with Activin A stimulation further substantiates the enhanced contractility of BMPR2^MutExDo^ iSMCs in comparison to WT cells, whereas less increase was detectable in BMPR2^MutKiDo^ iSMCs. This aligns with elevated pMLC levels in SMCs of some PAH patients ^33^. While αSMA, another component of the contractile apparatus was significantly upregulated in iSMCs for both mutations, Activin A stimulation led to significant upregulation only in BMPR2^MutExDo^ iSMCs in case of calponin 1. Supporting the findings of the contraction assay, pro-contractile genes are markedly upregulated in both BMPR2^Mut^-iSMC types with Activin A treatment compared to untreated cells. However, whether the observed differences in the expression of individual genes and MLC phosphorylation indicate mutation-specific effects remain speculative at this stage.

Interestingly, sotatercept co-application largely reversed the mutation-dependent increase in contractility induced by Activin A treatment in BMPR2^MutExDo^ iSMCs and completely reversed it in BMPR2^MutKiDo^ iSMCs. On the molecular level sotatercept significantly reduced expression of calponin-1 and αSMA in Activin A-stimulated BMPR2^MutExDo^ iSMCs, which is consistent with previous observations in pulmonary artery smooth muscle cells derived from PAH-patients (PASMCs) ^9^. In addition to structural effects, sotatercept reduced pMLC, which is linked to actomyosin contractile activity Together these findings suggest a hitherto unrecognized MoA of sotatercept, which may explain the rapid clinical response observed in a subgroup of patients, and which may operate synergistically with the known pro-apoptotic effects.

Another hallmark of PAH associated with pulmonary vascular remodeling is pathogenic deposition and composition of extracellular matrix in both distal and proximal pulmonary arteries ^5^. Dysregulation of ECM assembly occurs early in the pathogenesis, leading to increased vessel wall thickness, the formation of occlusive intimal lesions, the loss of vascular elasticity, and heightened vessel stiffness ^6^. While myofibroblasts appear during this process as the primary producers of ECM during development of PAH, it has been postulated that such cells may originate from local endothelial cells via EndoMT ^42^, or be derived from quiescent resident fibroblasts or bone marrow-derived circulating cells ^40,41^. Since myofibroblasts are known to produce large quantities of collagen I during pathological tissue remodeling and fibrosis ^67^, we hypothesized that the cells producing high levels of collagen I under Activin A treatment of BMPR2^Mut^ iSMCs, may actually represent myofibroblasts formed from SMCs via the process of smooth muscle cell – to – myofibroblast transition (SMC-MFT). This process has been demonstrated experimentally in atherosclerosis ^68^, but the contribution of SMC-MFT has not yet been investigated in HPAH.

Contractile properties, collagen I production and appearance of the characteristic myofibroblasts features were more pronounced in Activin A-treated BMPR2^MutExDo^ iSMCs and less prominent in BMPR2^MutKiDo^ iSMCs. This finding potentially indicates differential effects on downstream signal transduction pathways. The increased proliferation combined with the increased expression of collagen I and other genes associated with ECM formation, and with the upregulation of genes characteristic of myofibroblasts and other mesenchymal cell types in Activin A-treated BMPR2^Mut^ iSMCs, clearly indicates that alterations of the SMC phenotype in HPAH extends beyond the established contractile-to-synthetic switch ^65^ and indicates pathological transformation of BMPR2^MutExDo^ iSMCs into myofibroblasts. As our iSMC cultures consisted of highly enriched SMCs, and given that the upregulation of collagen I and αSMA filaments was observed in the vast majority of cells, it can be concluded that contaminating ECs or fibroblasts were not the source of the detected myofibroblasts. By contrast, the efficiency of myofibroblast formation following Activin A treatment suggests that SMC-MFT is the primary mechanism of myofibroblast formation in the arteries of PAH patients.

Notably, sotatercept treatment also significantly mitigated the effects of dysregulated ECM production and expression of myofibroblast-associated markers and prevented the conversion of SMCs into myofibroblasts which may further contribute to the drug’s anti-remodeling effects. Based on recently published data, we hypothesized that, in the myofibroblasts derived from our BMPR2^Mut^ iSMCs, mechanotransduction via ECM proteins and integrin complexes (ITGA/B) is creating a positive feedback loop ^59,69^ that promotes vascular remodeling in PAH ^61^. Integrin α2 and β1 subunits form heterodimers that can bind to collagen I and serve as transmembrane receptors for outside-in signaling, thereby enabling mechanotransduction and stimulating TGFβ release and - signaling. We detected upregulated ITGA levels exclusively in Activin A-stimulated BMPR2^MutExDo^-iSMCs, which represent an additional mediator in SMC related PAH pathogenesis. In PAH, so far only other types of integrins, such as α1, α8, αv and αVβ3, ^60,70^, α5β1 ^71^, β1 ^72^ have been described to play a role in PAH pathogenesis. The lack of integrin α2 upregulation in Activin A-stimulated BMPR2^MutKiDo^-iSMCs, which are less contractile than Activin A-stimulated BMPR2^MutExDo^-iSMCs, may reflect the typical downregulation of integrin α2 expression in contractile SMCs ^73^. Interestingly, we observed that sotatercept rescued the upregulation of integrin α2. Given that Activin A modulates collagen–integrin-mediated mechanotransduction and actomyosin contractility, both of which are closely linked to TGF-β signaling, we hypothesized that TGF-β pathway activity might be affected as well. We observed a mutation-dependent elevated expression of TGFβ receptor type I (ALK5) selectively in Activin A-stimulated BMPR2^MutExDo^ iSMCs, which is consistent with increased TGFβR1 levels reported in a subset of PAH lung tissues ^12^. Interestingly, this was suppressed by co-treatment with sotatercept.

Following the integration of our findings into the existing understanding of PAH pathomechanisms, we propose a comprehensive model of PAH induction and pathogenesis (Fig. 5):

In vascular SMCs of healthy individuals, Activin A via ACVR1B/ACVR2A/B tetramers ^74^, and TGFβ through TGFβR1/TGFβR2, all contribute to the activation of TGFβ signaling via SMAD2/3 ^75^. On the other hand, BMP10 homo- and BMP10/BMP9 heterodimers, present in plasma and tissue fluids ^76,77^, with both BMPs showing high affinity binding of ACVRL1, and BMP10 showing higher affinity binding of BMPR2^78^, effectively activate BMP signaling *in vivo* via ACVRL1/BMPR2 or ACVR1/BMPR2 and SMAD1/5/9 ^75^, which in turn inhibits SMAD2/3 phosphorylation via SMAD7, thus limiting TGFβ signaling ^79,80^. Moreover, the presence of sufficient BMPR2 protein on the cell membrane also prevents surplus formation of high-affinity receptor complexes for TGFβ signaling ^81^. These inhibitory mechanisms balance the positive feedback loop of the TGFβ pathway, which acts via an increased release of TGFβ from ECM stores mediated by mechanotransduction ^61,81^.

In SMCs of patients carrying heterozygous BMPR2^MutExDo^, reduction of SMAD 1/5/9 phosphorylation and in addition a shift to mixed hetereomeric receptor complexes that bind TGFβ significantly enhance SMAD2/3 phosphorylation. In case of the heterozygous BMPR2^MutKiDo^ mutations, the mutated BMPR2 still reaches the surface ^82^, but the mutation leads to reduction or abrogation of SMAD 1/5/9 phosphorylation and enhanced SMAD2/3 phosphorylation in SMCs.

A previous report suggested, that in patients with strong PAH phenotype carrying heterozygous BMPR2 mutations, a decreased expression of the BMPR2 WT allele may cause an additional shift to mixed hetereomeric receptor complexes that preferentially bind TGFβ^81^. However, we did not observe such decreased BMPR2 expression in our patient-derived iPSC-SMCs.

At this point we would like to highlight an important difference between the hypothesized *in vivo* scenario described above, and our *in vitro* iPSC-SMC model with reduced complexity. SMCs generally do not express BMP9 or BMP10. Accordingly, our RNA-Seq data also show no BMP9/10 expression in the iSMCs. Although our RNASeq data reveal that the cultured iSMCs do express some BMP4, binding of BMP2 and BMP4 does presumably not lead to significant phosphorylation of SMAD1/5/9, since nanomolar concentrations of both ligands are required for transient phosphorylation ^83,84^, while picomolar concentrations of BMP9/BMP10 result in long-lasting phosphorylation of SMAD 1/5/9 ^85^. Moreover, in contrast to BMP9/10, BMP2/4 mediated SMAD1/5/9 activation is inhibited by Noggin, which is expressed in the iSMCs ^83^. All this likely explains the lack of SMAD1/5/9 activation in all experimental groups including WT iSMCs, and iSMC with heterozygous BMPR2^MutExoDo^ or BMPR2^MutKiDo^ mutations.

In our model, sufficient BMPR2 on the surface of WT SMCs may limit the proportion of heteromeric receptor complexes with high affinity to TGFβ, thus preventing excessive SMAD2/3 phosphorylation. Binding of Activin A to ACVR1B/ACVR2A/B complexes may on one hand lead to additional SMAD2/3 activation, but internalization of these receptor complexes may further limit SMAD2/3 phosphorylation.

In contrast to WT cells, in our iSMCs carrying a heterozygous mutation in the BMPR2 extracellular domain, the mutation may not only prevent binding of BMP ligands, but may also prevent the receptor from reaching the cell surface or cause mRNA degradation ^86^. This should lead to a shift to mixed heteromeric receptor complexes that preferentially bind TGFβ and strongly enhance SMAD2/3 phosphorylation in SMCs thereby supporting the positive TGFβ feedback loop.

In SMCs carrying a heterozygous missense mutation (c.1471C>T, R491W) in exon 11 in the BMPR2 kinase domain, receptors reach the plasmamembrane ^82,87^, and a relevant shift to heteromeric mixed receptor tetramers that bind TGFβ can be excluded. Also, a decreased level of total BMPR2 expression from both alleles was not detected (Supplementary figure 8 A). Therefore, the observed increased SMAD2/3 phosphorylation after Activin A treatment may be promoted by non-canonical pathways. Impaired BMPR2 kinase function, including the R491W mutation investigated here was shown to shift the balance towards TGFβ-TAK1-MAPK signaling, resulting in increased p38 ^88,89^ as observed also in SMCs of patients with BMPR2 mutations ^90^. Moreover, Activin A induced signaling via ACVR1B/ ACVR2A/B was shown to increase ERK1/2 and p38 phosphorylation ^91^; both factors are involved in SMC contractility and collagen I synthesis ^91,92^.

Activation of both, p38 and ERK can increase phosphorylation of MLC in SMCs ^92^, thereby enhancing TGFβ release and supporting the positive TGFβ feedback loop.

Furthermore, an activation of predominantly non-SMAD by actin-myosin ^84^ and additionally by collagen-integrin-interactions ^85^ contributes to this positive feedback loop. The observed upregulation of the integrin receptor subunit α2 may further support this process. As a result, a self-sustaining imbalance between BMP and TGFβ signaling is induced, which reinforces the expression of Activin A ^86–88^, collagen, and actin–myosin components again ^89,90^, thereby driving the severe pathological progression in PAH ^91^.

Given that PAH-related BMPR2 mutations are heterozygous, all downstream effects of these mutations will result solely in quantitative alterations to signaling pathways rather than complete abrogation. However, these quantitative shifts are apparently sufficient to cause minor pathological deviations that contribute to disease development in the long term.

In vivo treatment with the Activin/BMP ligand trap sotatercept ^13^ appears to eliminate a significant portion – or at least a substantial part - of the available Activin A, potentially also BMP10 and to a lesser extend BMP9 from serum and extracellular tissue fluids ^76,77^. Apparently, the attenuation of Activin A by sotatercept seems sufficient to downregulate the pathological feedback loop, which is otherwise sustained via binding of ECM-released TGFβ to TGFβR1/TGFβR2 ^93^, expressed Activin to ACVR1B / ACVR2A/B ^91^, and BMP9 to via ACVRL1/ ACVR2 A/B ^94^.

In addition, as shown by Wang et al., BMP9 (released by the liver) and BMP10 (released by the right atrium) are directly involved in SMC contractility ^95^, reduction of available BMP10 and to a lesser extend BMP9 by sotatercept could therefore lead to reduced SMC contractility by limiting BMP9/10-dependent p38 signaling^92,96^. We hypothesize that decreased contractility and increased apoptosis in SMCs and SMC-derived myofibroblasts account for the rapid response to sotatercept in some patients, while the blockade of SMC-MFT and the reversal of the tissue remodeling leads to full therapeutic benefit.

## Conclusion

We present the first PAH *in vitro* model based on patient iPSC-derived SMCs rather than iPSC-derived ECs. This model recapitulates key phenotypes of pulmonary vascular SMCs associated with PAH, including hyperproliferation, reduced apoptosis and dysregulated ECM deposition. It also revealed previously unrecognized cellular disease mechanisms, such as SMC-MFT, which leads to the accumulation of myofibroblasts and the aberrant deposition of extracellular matrix. Using this platform, we also investigated the mechanisms of action of the activin signaling inhibitor sotatercept. We confirmed its antiproliferative and proapoptotic effects and demonstrated inhibition of SMC contractility and SMC-MFT, which likely contribute to its therapeutic effects.

## Supplementary Figure legends

**Supplementary figure 1:**
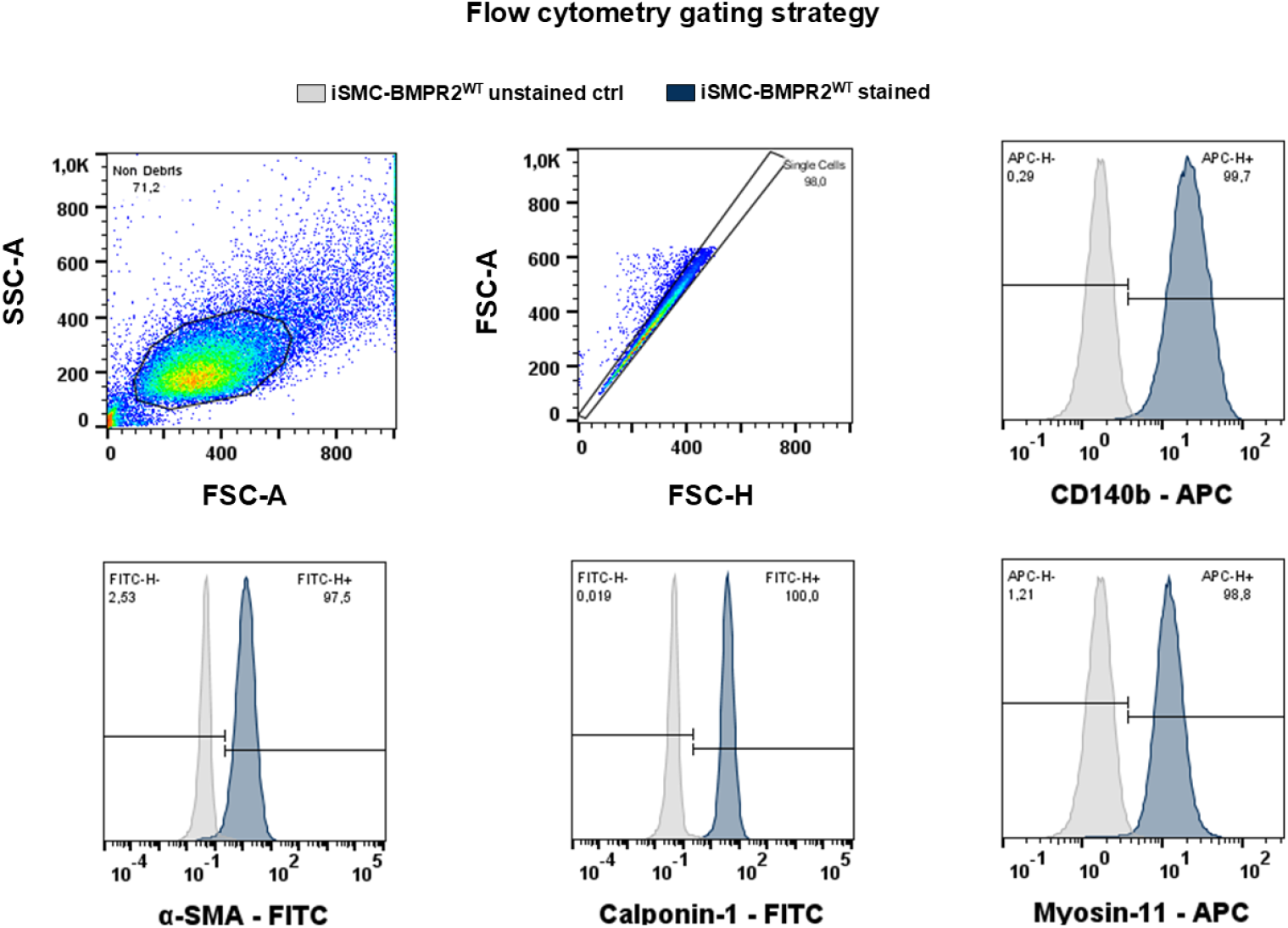
Flow-cytometric characterization of iSMCs. Representative gating strategy and SMC marker expression in BMPR2^WT^-iSMCs. Cells were gated by forward and side scatter to exclude debris, followed by FSC-A/FSC-H gating to select singlets. Histograms show staining for CD140b (PDGFRβ), α-smooth muscle actin (α-SMA), calponin-1 and myosin-11 (blue) relative to the unstained control (grey). Percentages indicate marker-positive cells, with gates defined using the unstained control. FSC-A, forward scatter-Area, FSC-H, forward scatter heigth; SSC-A, side scatter area; APC, allophycocyanin; FITC, fluorescein isothiocyanate.

**Supplementary figure 2:**
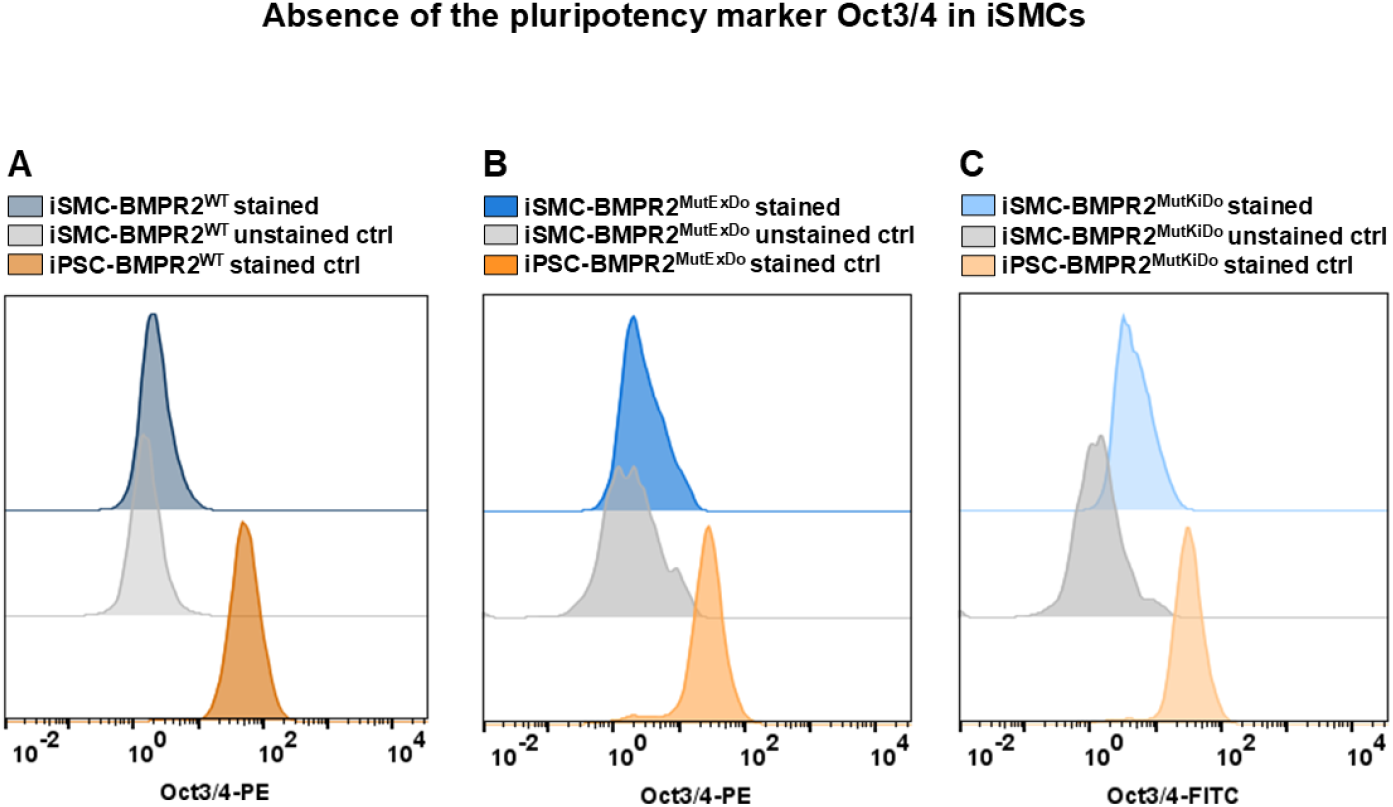
Absence of cells expressing the pluripotency marker OCT4 among iSMCs. Representative flow-cytometric analysis of OCT3/4 expression in iSMCs derived from (**A**) BMPR2^WT^, (**B**) BMPR2^MutExDo^ and (**C**) BMPR2^MutKiDo^. Stained iSMCs are shown in blue, genotype-matched unstained iSMC controls in grey, and the corresponding parental hiPSC positive controls in orange.

**Supplementary figure 3:**
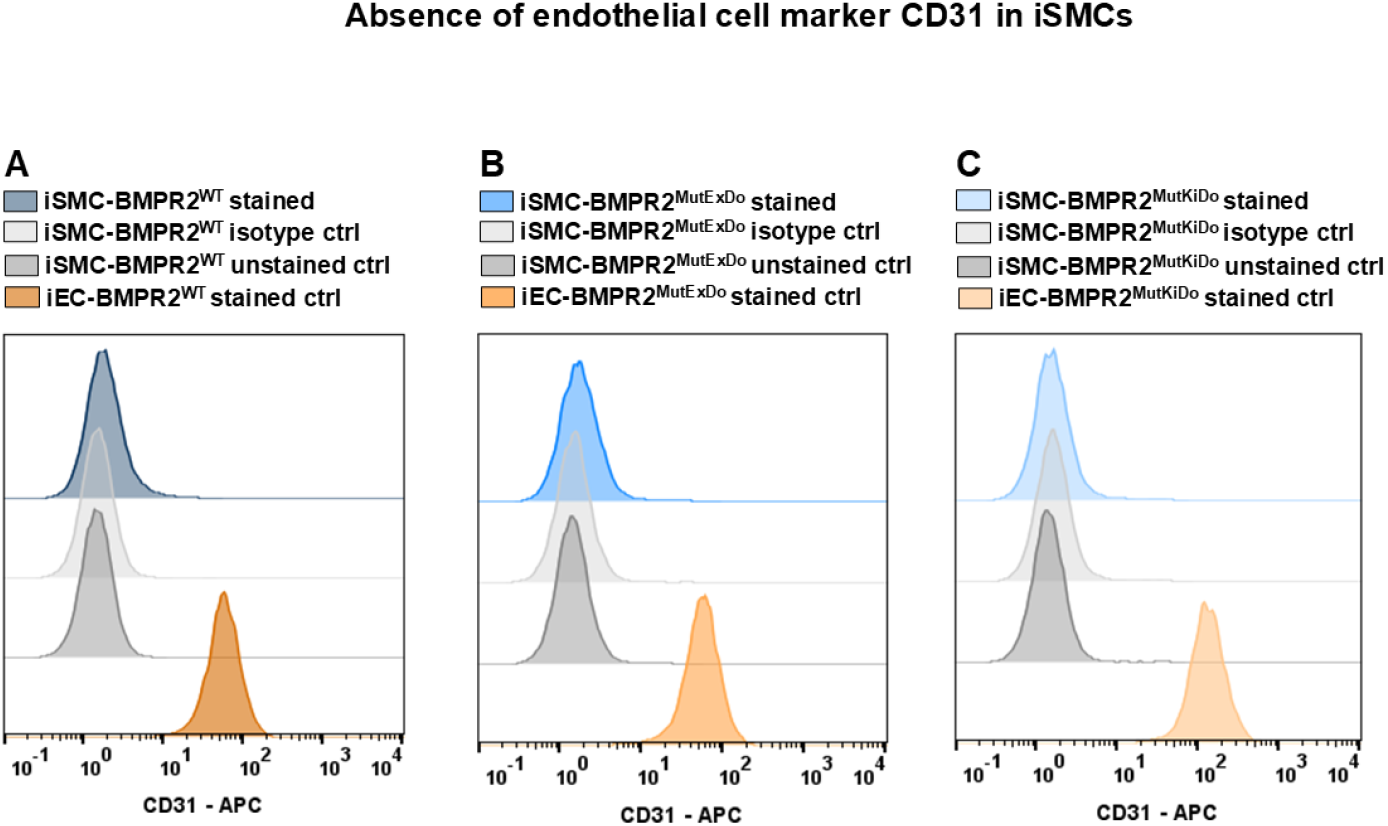
Absence of endothelial cell marker expression in iSMCs. Representative flow-cytometric analysis of CD31 expression in iSMCs derived from (**A**) BMPR2^WT^, (**B**) BMPR2^MutExDo^ and (**C**) BMPR2^MutKiDo^. Stained iSMCs are shown in blue, genotype-matched isotype control in light grey, unstained iSMC controls in dark grey and iPSC-derived endothelial cells as positive controls in orange.

**Supplementary figure 4:**
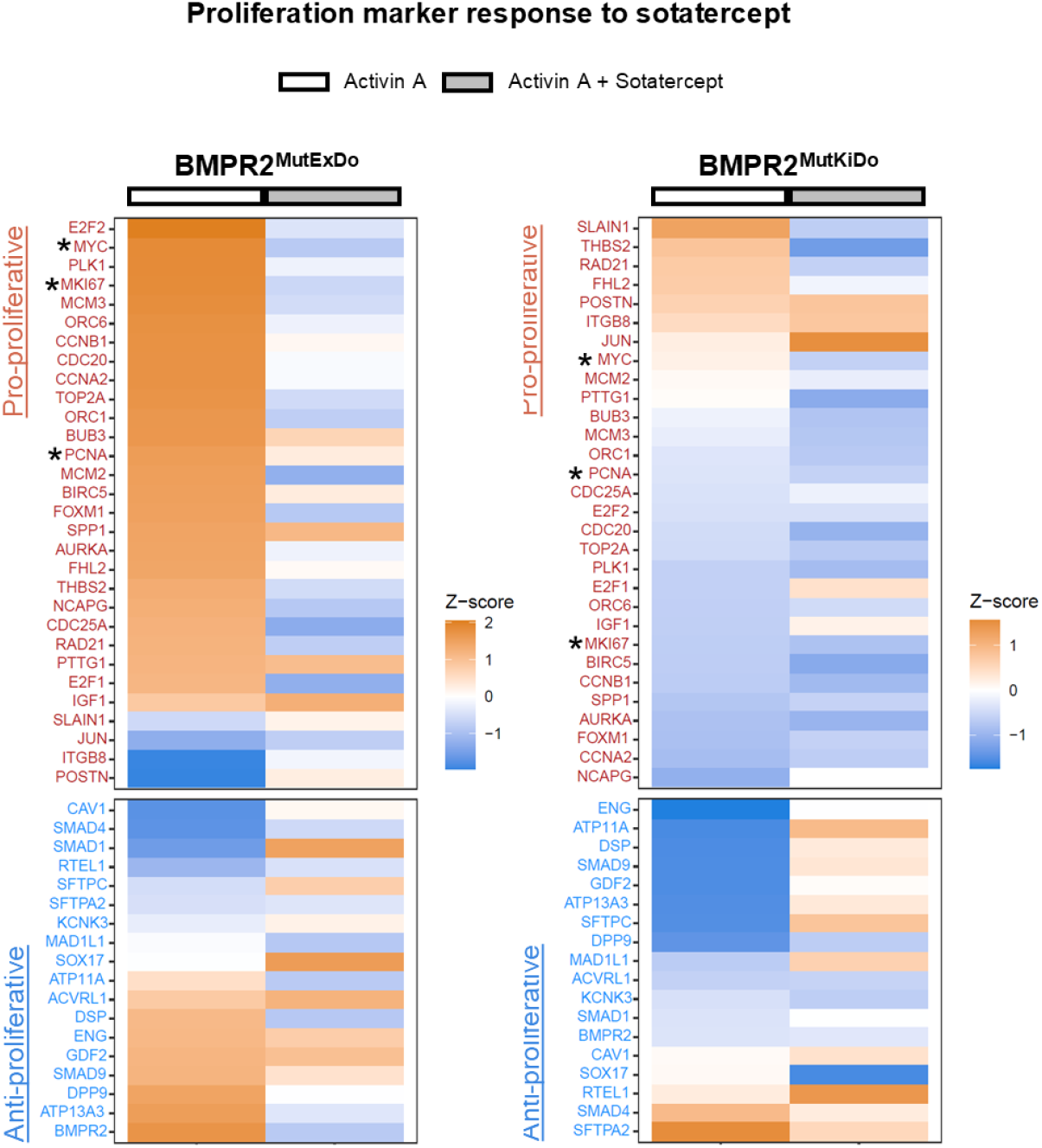
Sotatercept leads to downregulation of proproliferative genes in Activin A-treated BMPR2^mutExDo^ iSMCs. Heatmaps showing differentially expressed genes associated with proliferation in Activin A vs. Activin A + sotatercept treated iSMCs. For proliferation gene set (MSigDB: M16210) scoring, bulk RNAseq expression values were analyzed using a gene-by-sample FPKM matrix (mean FPKM of three independent biological replicates per group, Activin A and Activin A + sotatercept each condition normalized to the untreated control). Stars (*) highlight mentioned genes in the respective text.

**Supplementary figure 5:**
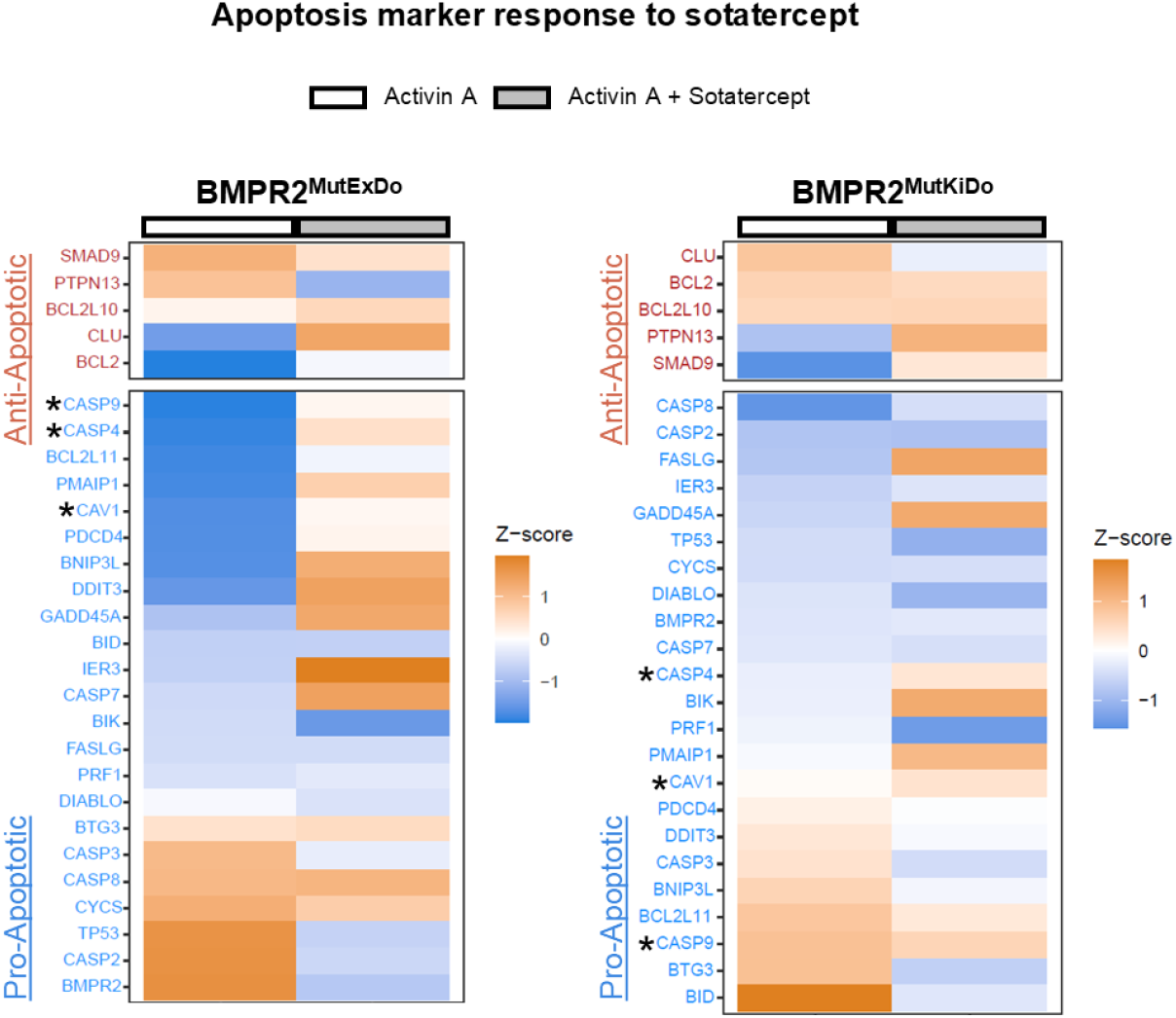
Sotatercept leads to upregulation of proapoptotic genes in Activin A-treated BMPR2^mutExDo^ iSMCs. Heatmaps showing differentially expressed genes associated with apoptosis in Activin A vs. Activin A + sotatercept treated iSMCs. For apoptosis gene set (MSigDB: M5902) scoring, bulk RNA-seq expression values were analyzed using a gene-by-sample FPKM matrix (mean FPKM of three independent biological replicates per group, Activin A and Activin A + sotatercept each condition normalized to the untreated control). Stars (*) highlight mentioned genes in the respective text.

**Supplementary figure 6:**
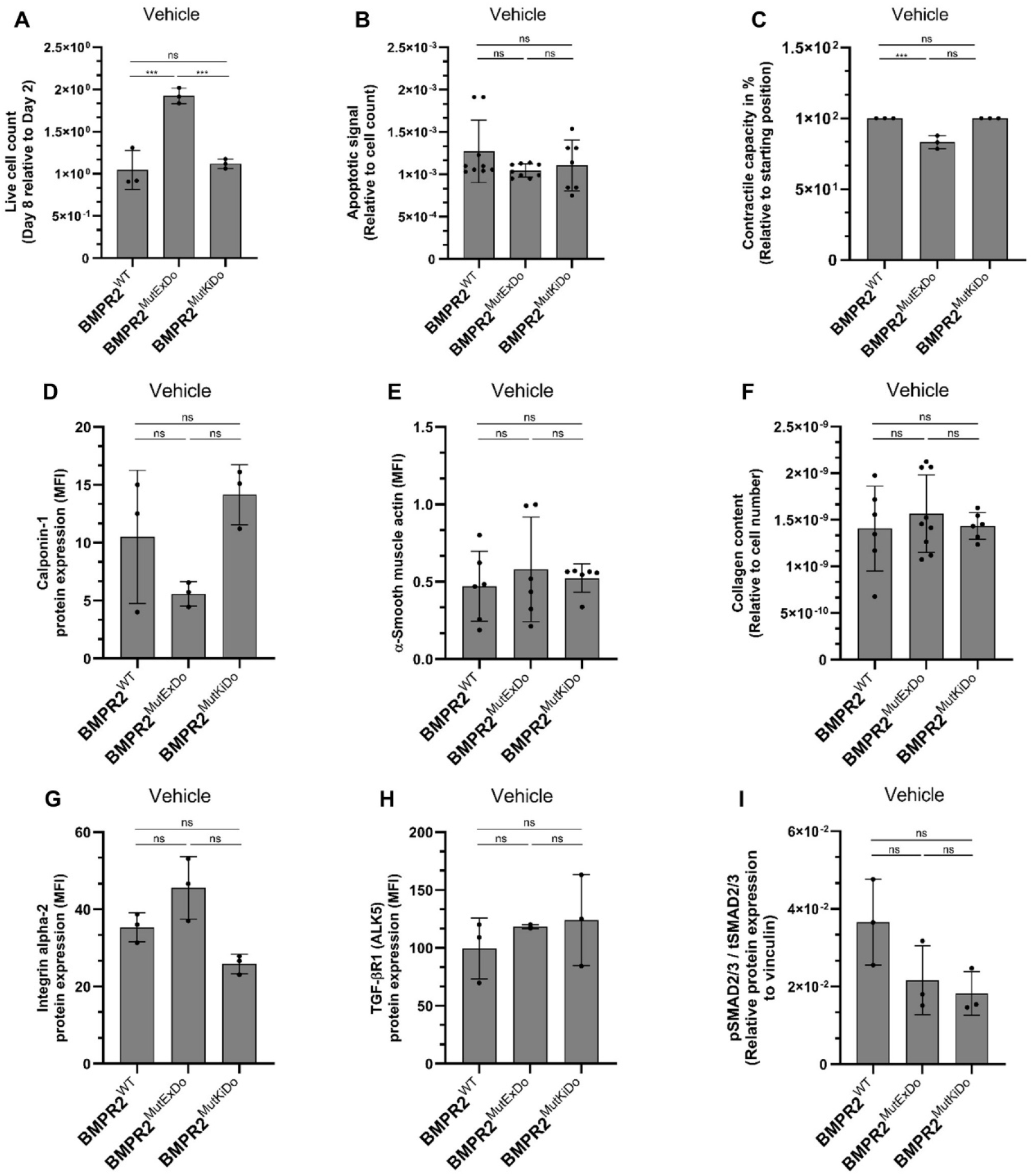
Baseline functional and molecular characterization of iSMCs. Vehicle-treated BMPR2^WT^, BMPR2^MutExDo^ and BMPR2^MutKiDo^ –iSMCs were assessed for (**A**) quantification of live cell numbers after 8 days in culture relative to day 2, (**B**) apoptotic signal on day 5 of cultivation relative to cell count by caspase activity, (**C**) contractile capacity in a collagen contraction assay, (**D**) calponin-1 expression by flow cytometry, (**E**) α-smooth muscle actin (α-SMA) expression by IF quantification, (**F**) collagen I deposition by SiriusRed assay, (**G**) integrin α2 expression, and (**H**) TGF-β receptor I (TGFBR1/ALK5) expression by flow cytometry, and (**I**) the ratio of phosphorylated to total SMAD2/3 western blot analysis. Data is presented as mean ± SD, n= 3 independent biological replicates per column indicated as dots. Statistical significance between genotypes was assessed using two-sided unpaired t-tests; ns, not significant; *p < 0.05, **p < 0.01, ***p < 0.001.

**Supplementary figure 7:**
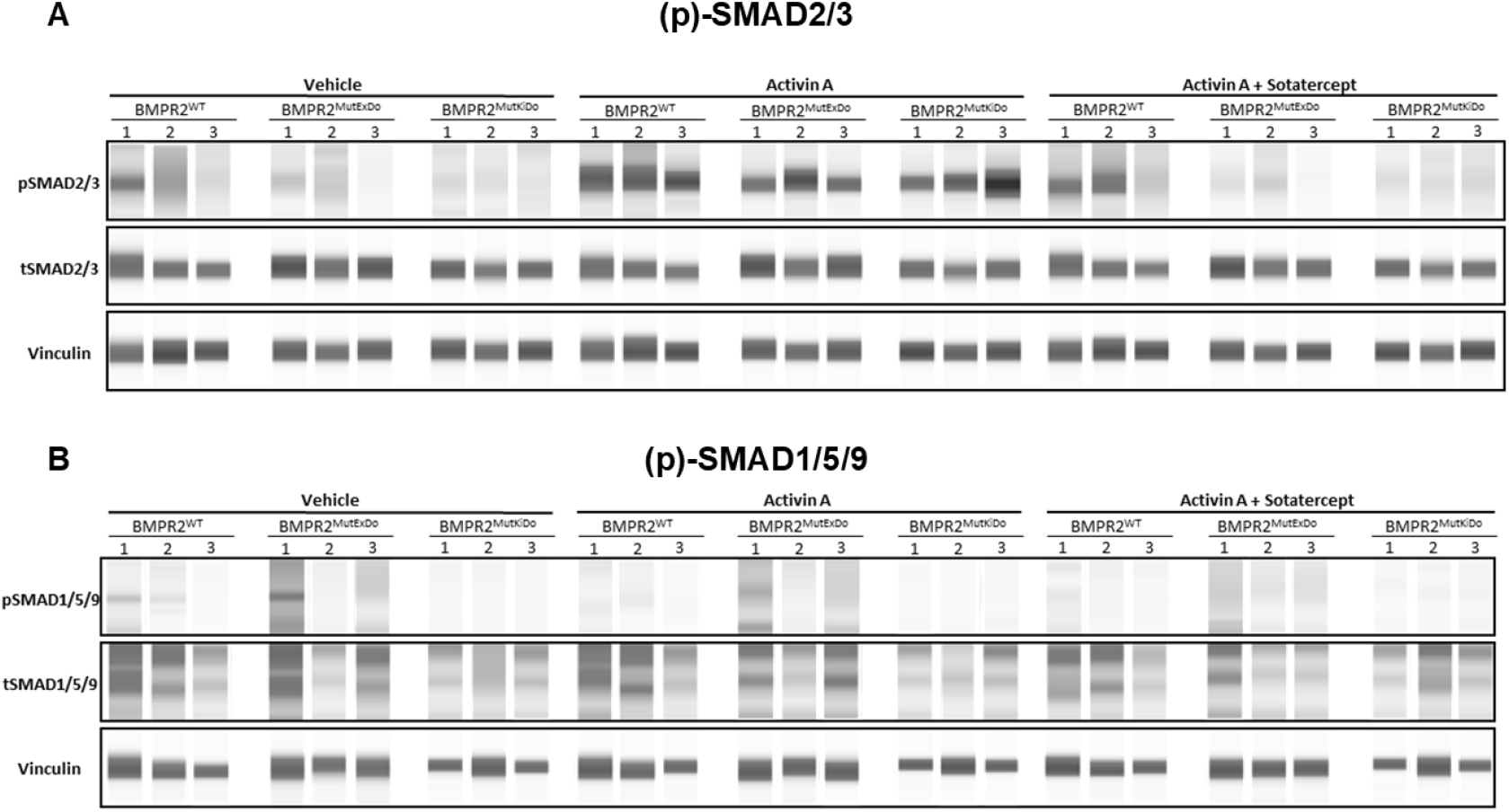
Protein expression analysis. **(A, B)** Virtual blot images from all biological replicates obtained by automated capillary-based western blot analysis of (A) p-SMAD2/3, t-SMAD2/3 (∼60 kDa) and vinculin (∼124 kDa), (B) p-SMAD1/5/9, t-SMAD1/5/9 (∼60 kDa) and vinculin (∼124 kDa).

**Supplementary figure 8:**
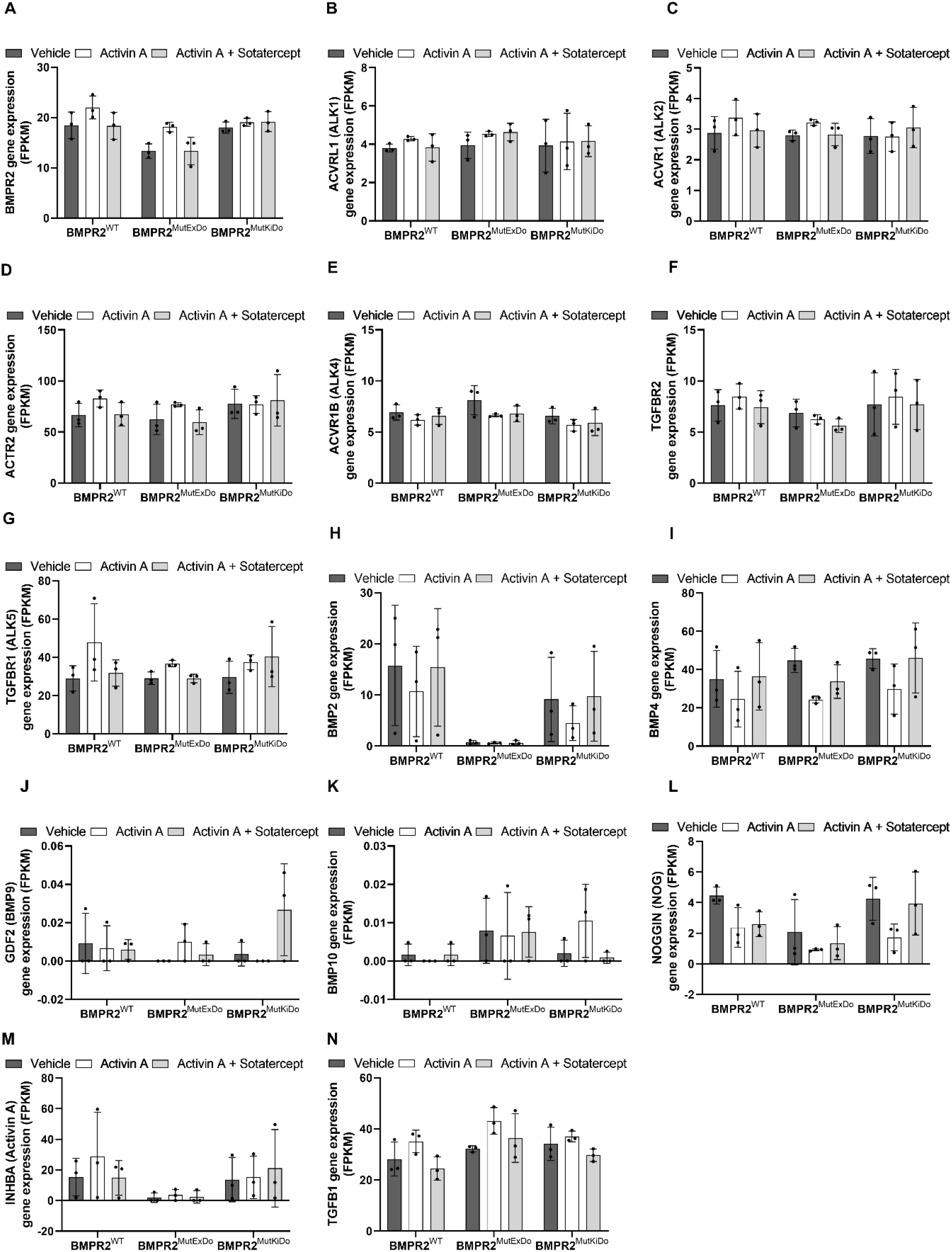
Gene expression analysis. Expression of TGF-β/BMP pathway receptors and ligands in BMPR2^WT^ and BMPR2-mutant iSMCs treated with vehicle, Activin A, or cotreated with Activin A and sotatercept. Shown are transcript levels of BMPR2, ACVR1, ACVR1B, ACVR2, ACVR2B, TGFBR1, TGFBR2, BMP2, BMP4, GDF2, BMP10, NOG, INHBA, and TGFB1, quantified by bulk RNA sequencing and expressed as FPKM.

## Material and Methods

### Human subjects

This study is an exploratory analysis to translate the results from the HPAH iSMC *in vitro* model to HPAH patients, to offer further insights into the therapeutic response to sotatercept in patients. For a paired analysis, TGFβ content in patient sera at baseline and on sotatercept treatment was examined from eight patients and on-treatment samples were obtained 9–41 months after sotatercept initiation. All patients signed the German Centre for Lung Research informed consent form (https://dzl.de/en/dzl-data-warehouse), which permits the use of clinical routine data and blood samples for research purposes. This form has been approved by our institution’s ethics committee (number 8540_BO_K_ 2019).”

### Sample collection and processing

Peripheral blood samples were taken from the same patients before and during therapy with sotatercept. Samples were collected in serum tubes, allowed to clot, processed within 2 h, and centrifuged at 1,500 g for 10 min at 4 °C. Serum was aliquoted and stored at -80 °C until ELISA analysis.

### ELISA-human TGFβ1 (Transforming growth factor beta-1)

Human transforming growth factor beta-1 (TGFβ1) was quantified in serum samples from HPAH patients by ELISA (Abcam, cat. no. ab100647). Latent TGFβ1 was activated by incubating 50 µL of 1 N HCL with 100 µL serum (FCS; PAN Biotech, cat. no. P30-3306) for 10 min at room temperature (RT). The acidified serum was neutralized by adding 50 µL of 1.2 N NaOH/0.5 M HEPES and assayed immediately. The sandwich ELISA was performed as follows: 100 µL of standard and sample was incubated overnight (ON) at 4°C with gentle shaking. The next day, wells were washed four times with 300 µL of the washing solution and plate was blotted against clean paper towels. In the next step, 100 µL of the 1x biotinylated TGFβ1 detection antibody per well was added and incubated for 1 h at RT with gentle shaking. Wells were washed again. Then, 100 µL of 1x HRP-streptavidin solution was added and incubated for 45 min at RT with gentle shaking. Wells were washed again and 100 µL of the TMB one-step substrate reagent was added and incubated for 30 min at RT in the dark with gentle shaking. Finally, 50 µL of the stop solution was added to each well and absorbance was measured at 450 nm.

### Human induced pluripotent stem cells (hiPSCs)

For HPAH *in vitro* modelling, three iPSC clones per patient cell line were compared to three independent cell lines from healthy individuals. The following human pluripotent stem cell lines were generated from CD34^pos^ cells using the CytoTune®-iPS 2.0 Sendai Reprogramming Kit (Thermo Fisher Scientific, cat. no. 16517), following the manufacturer’s instructions: WT1 (MHHi001-A; clone 2; male), WT2 (MHHi038-A; clone 20; male), WT3 (MHHi039-A clone 12; female), BMPR2^MutExDo^ heterozygous in-frame-deletion c.248-1 – 418+1 of exon 3 in the extracellular domain, (MHHi023-A; clone 18, 25, 26; female), BMPR2^MutKiDo^ (heterozygous missense mutation c.1471C>T in exon 11 in the kinase domain), MHHi024-A; clone 14, 16, 18; female). Pluripotency and genomic stability were tested for all hiPSC lines (Usman et al., 2021). Informed consent for blood collection, iPSC generation and anonymous data storage was obtained from the patients or legal caregivers according to the principles expressed in the Declaration of Helsinki (pediatric IRB approval #2200 to G.H., Hannover Medical School; 2926-2015). hiPSCs were maintained on vitronectin-coated (Gibco, cat. no. A31804) plates (Nunc, cat. no. 140675) in Essential 8 medium (DMEM/F12 (ThermoFisher, cat. no. 11330-057), 64 mg/L ascorbic acid 2-phosphate (Sigma-Aldrich, cat. no. A92902), 14 μg/L sodium selenite (Sigma-Aldrich, cat. no. 214485), 543 mg/L sodium bicarbonate (Sigma Aldrich, cat. No. 55761), 20 μg/L Insulin (Sigma-Aldrich, cat. no. I9278), 10.7 μg/mL human recombinant transferrin (Sigma-Aldrich, cat. no. T3705), 100 ng/mL bFGF (Peprotech, cat. no. AF-100-18B), 2 ng/mL TGFb1 (PeproTech, cat. no. 100-21C)). Cells were passaged every 3–4 days using Accutase^TM^ (Gibco, cat. no. A1110501) at 32,000 cells/cm^2^ and cultivated at 37 °C, 5% CO2. Cells were tested monthly for mycoplasma contamination using the MycoStrip® - Mycoplasma Detection Kit (Invivogen, cat. no. REP-MYS). Passages between 4 to 8 were used for starting directed differentiation into vascular smooth muscle cells.

### Differentiation in iPSC-derived smooth muscle cells (iSMCs)

Differentiation of iPSCs into vascular smooth muscle cells (iSMCs) was performed following a modified protocol derived from (Patsch et. al, 2015), described in detail below. On day 0, iPSCs were seeded at 12,000 cells/cm² and cultivated in Essential 8 medium (DMEM/F12 (ThermoFisher, cat. no. 11330-057), 64 mg/L ascorbic acid 2-phosphate (Sigma-Aldrich, cat. no. A92902), 14 μg/L sodium selenite (Sigma-Aldrich, cat. no. 214485), 543 mg/L sodium bicarbonate (Sigma Aldrich, cat. No. 55761), 20 μg/L Insulin (Sigma-Aldrich, cat. no. I9278), 10.7 μg/mL human recombinant transferrin (Sigma-Aldrich, cat. no. T3705), 100 ng/mL bFGF (Peprotech, cat. no. AF-100-18B), 2 ng/mL TGFb1 (PeproTech, cat. no. 100-21C)) with 10 µM Y-27632 (Tocris, cat. no. 1254) for 24 h. On day 1, medium was changed to Essential 6 medium (DMEM/F12 (ThermoFisher, cat. no. 11330-057), 64 mg/L ascorbic acid 2-phosphate (Sigma-Aldrich, cat. no. A92902), 14 μg/L sodium selenite (Sigma-Aldrich, cat. no. 214485), 543 mg/L sodium bicarbonate (Sigma Aldrich, cat. No. 55761), 20 μg/L Insulin (Sigma-Aldrich, cat. no. I9278), 10.7 μg/mL human recombinant transferrin (Sigma-Aldrich, cat. no. T3705) supplemented with 7.5 µM CHIR66021 (provided by the Institute of Organic Chemistry, Leibniz University, Hannover, Germany) and 25 ng/mL BMP4 (R&D Systems, cat. no. 314-BP) for mesoderm induction via Wnt modulation. On day 4, medium was replaced by Essential 6 medium supplemented with 2 ng/mL Activin A (Peprotech, cat. no. 120-14E) and 10 ng/mL PDGF-BB (Peprotech, cat. no. 100-14B) for vascular SMC specification. On day 6, the differentiation process was terminated by dissociating and seeding the cells on different formats for further analysis. The differentiation efficiency and therefore purity of generated iSMCs was assessed via CD140b in flow cytometry.

### Flow cytometric analysis

Flow analysis was used for detection of surface and intracellular markers of iSMCs. For assessing the differentiation efficacy, freshly differentiated iSMCs were stained with directly labeled CD140b-APC (APC, Miltenyi Biotec, cat. no. 130-121-052, RRID:AB_2783952, recombinant human, 1:50) in PBS pH7.4 (1X) (Gibco, cat. no. 70011-036) supplemented with 1 % FCS (PAN Biotech, cat. no. P30-3306) and 1 mM EDTA (Sigma Aldrich, cat. no. E5134) for 45 min and propidium iodide (Miltenyi Biotec, cat. no. 130-093-233, 1:100) for 5 min, to exclude dead cells. For the Oct3/4 and CD31 staining, iSMCs were fixed and permeabilized with 4% PFA and 90% MeOH directly after differentiation and stained with directly-conjugated anti-Oct3/4-PE (Miltenyi, 130-105-554, RRID:AB_2653085, 1:50) or CD31-APC (Miltenyi,130-092-652, RRID:AB_871661, 1:50). For the quantification of HPAH phenotypic markers, freshly differentiated iSMCs were seeded as passage 1 at 20.000 cells/cm^2^ on vitronectin-coated (Gibco, cat. no. A31804) plates (Nunc, cat. no. 140675) in 200 µL/cm^2^ Essential 6 (DMEM/F12 (ThermoFisher, cat. no. 11330-057), 64 mg/L ascorbic acid 2-phosphate (Sigma-Aldrich, cat. no. A92902), 14 μg/L sodium selenite (Sigma-Aldrich, cat. no. 214485), 543 mg/L sodium bicarbonate (Sigma Aldrich, cat. No. 55761), 20 μg/L Insulin (Sigma-Aldrich, cat. no. I9278), 10.7 μg/mL human recombinant transferrin (Sigma-Aldrich, cat. no. T3705) with 10 µM Y-27632 (Tocris, cat. no. 1254) and cultivated at 37°C and 5% CO_2_ for 24 h. Between day 1-12, every 48 h media was changed to 200 µL/cm^2^ E6 with additives of Activin A (Peprotech, cat. no. GMP120-14E, 0.25 ng/ml), sotatercept (MedChemExpress, HY-P99590, 10 µg/ml) or Activin A with sotatercept. On day 12, for the quantification of HPAH phenotypic surface markers cells were fixed with 4% paraformaldehyde (Sigma-Aldrich, cat. no.158127) for 10 min, with ice-cold 90% methanol (JT Baker, cat. no. 8045) for 15 min and stained with a primary antibody for TGFβR1 (Thermo Fisher Scientific, cat. no. PA5-142789, RRID:AB_2933432, rabbit, 1:100) in PBS pH7.4 (1X) (Gibco, cat. no. 70011-036) supplemented with 1 % FCS (PAN Biotech, cat. no. P30-3306) and 1 mM EDTA (Sigma Aldrich, cat. no. E5134) over night at 4 °C. Cells were stained with a secondary antibody anti-rabbit (Cy5, Jackson ImmunoResearch Labs, cat. no. 711-175-152, RRID:AB_2340607, donkey, 1:300) in PBS pH7.4 (1X) (Gibco, cat. no. 70011-036) supplemented with 1 % FCS (PAN Biotech, cat. no. P30-3306) and 1 mM EDTA (Sigma Aldrich, cat. no. E5134) for 1 h at RT. On day 12, for quantification of HPAH phenotypic intracellular or transmembrane markers, cells were fixed with 4% paraformaldehyde (Sigma-Aldrich, cat. no.158127) for 10 min, ice-cold 90% methanol (JT Baker, cat. no. 8045) for 15 min, permeabilized with 1% tritonX-100 (Sigma-Aldrich, cat. no. 93443) PBS pH7.4 (1X) (Gibco, cat. no. 70011-036) supplemented with 1 % BSA (Sigma-Aldrich, cat. no. A9418) for 20 min and stained with the primary antibody CNN1 (Santa Cruz Biotechnology, cat. no. sc-53136, RRID:AB_793529, mouse, 1:50) or integrin α2 (Abcam, cat. no. ab181548, RRID:AB_2847852, rabbit, 1:100) in PBS pH7.4 (1X) (Gibco, cat. no. 70011-036) supplemented with 1 % BSA (Sigma-Aldrich, cat. no. A9418) and 1% tritonX-100 (Sigma-Aldrich, cat. no. 93443) over night at 4 °C. Cells were stained with a secondary antibody anti-rabbit (Cy5, Jackson ImmunoResearch Labs, cat. no. 711-175-152, RRID:AB_2340607, donkey, 1:300) or anti-mouse (AF488, Jackson ImmunoResearch Labs, cat. no. 715-545-151, RRID:AB_2341099, donkey, 1:300) in PBS pH7.4 (1X) (Gibco, cat. no. 70011-036) supplemented with 1 % BSA (Sigma-Aldrich, cat. no. A9418) and 1 % tritonX-100 (Sigma-Aldrich, cat. no. 93443) for 1 h at RT. Flow analysis was performed with the MACSQuant Analyzer 10 and data processing occurred with FlowJo_v10.8.1.

### Immunofluorescence staining

Immunofluorescence (IF) staining was performed to visualize characteristic markers of SMCs and HPAH phenotypic markers. For the quantification of HPAH phenotypic markers, freshly differentiated iSMCs were seeded as passage 1 at 7.000 cells/cm^2^ on vitronectin-coated (Gibco, cat. no. A31804) chamber slides in 350 µL/cm^2^ Essential 6 (DMEM/F12 (ThermoFisher, cat. no. 11330-057), 64 mg/L ascorbic acid 2-phosphate (Sigma-Aldrich, cat. no. A92902), 14 μg/L sodium selenite (Sigma-Aldrich, cat. no. 214485), 543 mg/L sodium bicarbonate (Sigma Aldrich, cat. No. 55761), 20 μg/L Insulin (Sigma-Aldrich, cat. no. I9278), 10.7 μg/mL human recombinant transferrin (Sigma-Aldrich, cat. no. T3705)) with 10 µM Y-27632 (Tocris, cat. no. 1254) and cultivated at 37°C and 5% CO_2_ for 24 h. Between day 1-12, every 48 h media was changed to 200 µL/cm^2^ E6 with additives of Activin A (Peprotech, cat. no. GMP120-14E, 0.25 ng/ml), sotatercept (MedChemExpress, HY-P99590, 10 µg/ml) or Activin A with sotatercept. On day 12, for the stainings of actin, CNN1 and TGFβR1 cells were fixed with 4% paraformaldehyde Sigma-Aldrich, cat. no.158127) for 10 min at RT, blocked with 10 % donkey serum (PAN Biotech, cat. no P30-0102) 10 min at RT and stained for actin (Santa Cruz Biotechnology, cat. no. sc-130616, RRID:AB_1561784, mouse, 1:50), CNN1 (Santa Cruz Biotechnology, cat. no. sc-53136, RRID:AB_793529, mouse, 1:50) or TGFβR1 (Thermo Fisher Scientific, cat. no. PA5-142789, RRID:AB_2933432, rabbit, 1:50) ON at 4°C. Cells were stained with a secondary antibody anti-rabbit (Cy5, Jackson ImmunoResearch Labs, cat. no. 711-175-152, RRID:AB_2340607, donkey, 1:300) or anti-mouse (AF488, Jackson ImmunoResearch Labs, cat. no. 715-545-151, RRID:AB_2341099, donkey, 1:300) for 1 h at RT. Stainings for MYH11, COL1: Cells were fixed with 4% paraformaldehyde (Sigma-Aldrich, cat. no.158127) for 10 min at RT, treated with citrate buffer for 10 min at 95 °C, blocked with 10 % donkey serum (PAN Biotech, cat. no P30-0102) for 10 min at room temperature and stained with MYH11 (Abcam, cat. no. ab133567, RRID:AB_2890982, rabbit, 1:50) or COL1 (Sigma-Aldrich, cat. no. C2456, RRID:AB 476836, mouse, 1:500) ON at 4°C. Cells were stained with a secondary antibody anti-rabbit (Cy5, Jackson ImmunoResearch Labs, cat. no. 711-175-152, RRID:AB_2340607, donkey, 1:300) or anti-mouse ((AF 488, Jackson ImmunoResearch, cat. no. 715-545-151, RRID:AB_2341099, donkey, 1:300) for 1 h at RT. Stainings for integrin α2: Cells were fixed with 4% paraformaldehyde (Sigma-Aldrich, cat. no.158127) for 10 min, ice-cold 90% methanol (JT Baker, cat. no. 8045) for 15 min and permeabilized with 1% tritonX-100 (Sigma-Aldrich, cat. no. 93443) in PBS pH7.4 (1X) (Gibco, cat. no. 70011-036) supplemented with 1 % BSA (Sigma-Aldrich, cat. no. A9418) for 20 min followed by blocking with 10 % donkey serum (PAN Biotech, cat. no P30-0102) for 10 min at RT and stained for integrin α2 (Abcam, cat. no., ab181548, RRID:AB_2847852, rabbit, 1:100) in TBS (pH 7.6, Tris 1X, in-house made) supplemented with 0.2% tritonX-100 (Sigma-Aldrich, cat. no. 93443) and 1 % BSA (Sigma-Aldrich, cat. no. A9418) ON at 4 °C. Cells were stained with a secondary antibody anti-rabbit (Cy5, Jackson ImmunoResearch Labs, cat. no. 711-175-152, RRID:AB_2340607, donkey, 1:300) in 0.2% tritonX-100 (Sigma-Aldrich, cat. no. 93443) in TBS (pH 7.6, Tris 1X, in-house made) supplemented with 1 % BSA (Sigma-Aldrich, cat. no. A9418) for 1 h at RT. All stainings were co-stained for nuclei detection with 0.167 µg/ml DAPI (Sigma Aldrich, cat. no. D9541) for 10 min at RT. IF images were taken with the Zen 2.6 software at the Axioscope 7 microscope (ZEISS) and equally detected in terms of laser intensity, exposure time and sub-sequential image processing.

### Western blots

Quantification of total and phosphorylated proteins of interests was performed by western blot analysis. On day 0, freshly differentiated iSMCs were seeded as passage 1 at 20.000 cells/cm^2^ on vitronectin-coated (Gibco, cat. no. A31804) plates (Nunc, cat. no. 140675) in 200 µL/cm^2^ Essential 6 (DMEM/F12 (ThermoFisher, cat. no. 11330-057), 64 mg/L ascorbic acid 2-phosphate (Sigma-Aldrich, cat. no. A92902), 14 μg/L sodium selenite (Sigma-Aldrich, cat. no. 214485), 543 mg/L sodium bicarbonate (Sigma Aldrich, cat. No. 55761), 20 μg/L Insulin (Sigma-Aldrich, cat. no. I9278), 10.7 μg/mL human recombinant transferrin (Sigma-Aldrich, cat. no. T3705)) with 10 µM Y-27632 (Tocris, cat. no. 1254) and cultivated at 37°C and 5% CO_2_ for 24 h. Between day 1-5, every 48 h media was changed to E6 with 200 µL/cm^2^. On day 6, media was changed to 200 µL/cm^2^ E6 with additives for 1 h: Activin A (Peprotech, cat. no. GMP120-14E, 25 ng/ml), sotatercept (MedChemExpress, HY-P99590, 466 ng/ml). Cells were lyzed and total protein was extracted with (Sigma Aldrich, cat. no. R0278) and protease-phosphatase-inhibitor (Thermo Fisher, cat. no. 78440). After centrifugation at 14.000 g for 20 min at 4°C supernatant was collected and stored at -80°C. The following primary antibodies were used: Vinculin (Sigma-Aldrich,cat. no. V9131, RRID:AB_477629, mouse, 1:125.000), total myosin light chain (Cell Signaling Technology, cat. no. 8505, RRID:AB_2728760, rabbit, 1:20), phosphorylated myosin light chain (Cell Signaling Technology, cat. no. 3671, RRID:AB_330248, rabbit, 1:50), phosphorylated Smad1/5/8 (Cell Signaling Technology, 13820, RRID:AB_2493181, rabbit, 1:20), total Smad1/5/8 ((Abcam, ab80255, RRID:AB 1658979, rabbit, 1:10), phosphorylated Smad2/3 (Cell Signaling Technology, 8828, RRID:AB_2631089, rabbit.,1:20) and total Smad2/3 (Cell Signaling Technology, 8685, RRID:AB_10889933, rabbit, 1:250)., secondary anti-rabbit (Biotechne, DM-001, RRID:AB_3095297, goat, ready-to-use) and secondary anti-mouse (Biotechne, DM-002, RRID:AB_3095298, goat, ready-to-use) Western blots were conducted with an automated western blot instrument, Jess ProteinSimple^TM^. The Jess Simple Western system (ProteinSimple, San Jose CA, USA) is an automated, capillary-based immunoassay enabling size-based separation and detection of proteins of interest in low µg quantities. According to the manufacturer’s instructions, 2.3 µL of protein lysate was mixed with 0.6 µL of 5x fluorescence master mix and 1 µL 0,1x sample buffer supplied in the standard reagent pack. After denaturation at 95°C for 5 minutes, samples were kept on ice and loaded onto the assay plate together with all required reagents (i.e., antibodies, wash and stripping/replex buffer, luminol/peroxide), followed by centrifugation for 5 minutes at 1,000 *g.* The assay plate and a 12–230 kDa capillary cartridge were loaded into the Jess instrument, where automated separation followed by a chemiluminescence-based immunodetection was completed within 5 hours. Chemiluminescent signals were detected using a CCD camera, and protein expression levels reflected by band intensities were visualized as electropherograms. Densitometric quantification was performed by calculating the area under the curve for each protein of interest using the Compass software (version 7.0.0, ProteinSimple). Normalized expression ratios were calculated as specified in the figure legends.

### Collagen assay

Quantification of collagen production was performed by a SiriusRed-based assay. On day 0, freshly differentiated iSMCs were seeded as passage 1 at 15.625 cells/cm^2^ on vitronectin-coated (Gibco, cat. no. A31804) plates (Nunc, cat. no. 140675) in 390 µL/cm^2^ Essential 6 Essential 6 (DMEM/F12 (ThermoFisher, cat. no. 11330-057), 64 mg/L ascorbic acid 2-phosphate (Sigma-Aldrich, cat. no. A92902), 14 μg/L sodium selenite (Sigma-Aldrich, cat. no. 214485), 543 mg/L sodium bicarbonate (Sigma Aldrich, cat. No. 55761), 20 μg/L Insulin (Sigma-Aldrich, cat. no. I9278), 10.7 μg/mL human recombinant transferrin (Sigma-Aldrich, cat. no. T3705)) with 10 µM Y-27632 (Tocris, cat. no. 1254) and cultivated at 37°C and 5% CO_2_ for 24 h. Between day 1 to 5, every 48 h media was changed to 390 µL/cm^2^ E6 with additives: Activin A (Peprotech, cat. no. GMP120-14E, 0.25 ng/ml), sotatercept (MedChemExpress, HY-P99590, 10 µg/ml). On day 6, collagen assay was performed as follows: Cells were incubated with 1 µg/ml Hoechst33342 (Thermo Fisher, cat. no. H3570) at 37°C and 5% CO_2_ for 1 h and emission was read at 465 nm (9×9 area scan, bottom) with a plate reader for nuclei counterstain. Cells were fixed with 4% paraformaldehyde (Sigma-Aldrich, cat. no.158127) for 10 min at RT and washed with DPBS (Gibco, cat. no. 14040-091). Cells were stained with SiriusRed (0.1% DirectRed 80 (Sigma Aldrich, cat. no. 365548) in 1% picric acid (Sigma Aldrich, cat. no. P6744)) for 30 min at RT. After three washing steps with HCL (0.01 M), bound SiriusRed was eluated with NaOH (0.1 M). Absorbance was measured at 570 nm with the multimode plate reader (Paradigm, Molecular devices), which is proportional to the amount of fibril collagen.

### Proliferation assay

To assess the proliferative capacities a MTT assay was conducted. On day 0, freshly differentiated iSMCs were seeded as passage 1 at 31.250 cells/cm^2^ on vitronectin-coated (Gibco, cat. no. A31804) plates (Nunc, cat. no. 140675) in 390 µL/cm^2^ Essential 6 (DMEM/F12 (ThermoFisher, cat. no. 11330-057), 64 mg/L ascorbic acid 2-phosphate (Sigma-Aldrich, cat. no. A92902), 14 μg/L sodium selenite (Sigma-Aldrich, cat. no. 214485), 543 mg/L sodium bicarbonate (Sigma Aldrich, cat. No. 55761), 20 μg/L Insulin (Sigma-Aldrich, cat. no. I9278), 10.7 μg/mL human recombinant transferrin (Sigma-Aldrich, cat. no. T3705)) with 10 µM Y-27632 (Tocris, cat. no. 1254) and cultivated at 37°C and 5% CO_2_ for 24 h. Between day 1 to 7, every 48 h media was changed to 370 µL/cm^2^ E6 with additives: Activin A (Peprotech, cat. no. GMP120-14E, 0.25 ng/ml), sotatercept (MedChemExpress, HY-P99590, 10 µg/ml). On day 8, proliferation assay was performed with the MTT cell proliferation kit (Abcam, cat. no. Ab211091). Cells were treated with 100 µL of a 1:1 mix of the MTT reagent in E6 per well and incubated at 37°C and 5% CO_2_ for 3 h. Subsequently, 150 µL of the MTT solvent were added to the MTT reagent in E6 per well and incubated on an orbital shaker at 37°C and 5% CO_2_ for 15 min. Solution was gently mixed and absorbance was measured at 590 nm. Values of additive treated wells were normalized to only E6 treated cells per cell line and clone.

### Apoptosis assay

The apoptotic rate was analyzed by a caspase-3/7 red assay. On day 0, freshly differentiated iSMCs were seeded as passage 1 at 31.250 cells/cm^2^ on vitronectin-coated (Gibco, cat. no. A31804) plates (Nunc, cat. no. 140675) in 390 µL/cm^2^ Essential 6 (DMEM/F12 (ThermoFisher, cat. no. 11330-057), 64 mg/L ascorbic acid 2-phosphate (Sigma-Aldrich, cat. no. A92902), 14 μg/L sodium selenite (Sigma-Aldrich, cat. no. 214485), 543 mg/L sodium bicarbonate (Sigma Aldrich, cat. No. 55761), 20 μg/L Insulin (Sigma-Aldrich, cat. no. I9278), 10.7 μg/mL human recombinant transferrin (Sigma-Aldrich, cat. no. T3705)) with 10 µM Y-27632 (Tocris, cat. no. 1254) and cultivated at 37°C and 5% CO_2_ for 24 h. On day 1 and 3, media was changed to 370 µL/cm^2^ E6 with additives: Activin A (Peprotech, cat. no. GMP120-14E, 0.25 ng/ml), sotatercept (MedChemExpress, HY-P99590, 10 µg/ml). On day 5, caspase staining was performed with the kit CellEvent™ Caspase-3/7 (ThermoFisher, cat. no. C10431). A 10x working caspase reagent mix was freshly prepared by 1:10 dilution of the 100x stock, combined with 1:10 of HOECHST33342 (Thermo Fisher, cat. no. H3570) for nuclear counterstaining in E6. Then, 14 µL of this 10x mix were added to the preexisting medium of 125 µl per well, leading to an end concentration of 1 µg/ml of HOECHSt33342. Staining mix was gently mixed and incubated for 60 min at 37 °C, 5% CO_2_. Fluorescence intensity was measured using a multimode plate reader (Paradigm, Molecular devices). Measurements were performed from the bottom of the 96-well plates to ensure optimal detection of cell-associated signals. For the caspase-3/7 red assay, excitation was set at 585 nm, and emission was recorded at 635 nm with an integration time of 140 ms in monochromatic mode. For nuclear counterstaining with HOECHST33342 (Thermo Fisher, cat. no. H3570), excitation was performed at 360 nm and emission was collected at 465 nm using the same integration time (140 ms) and monochromatic mode. To improve measurement accuracy, both caspase and HOECHST33342 signals were acquired using an AreaScan protocol (9 × 9 reads per well), with HOECHST33342 serving as the nuclear counterstain for normalization.

### Contraction assay

The contractility was assessed by a functional collagen-contraction assay. On day 0, freshly differentiated iSMCs were seeded as passage 1 at 20.000 cells/cm^2^ on vitronectin-coated (Gibco, cat. no. A31804) plates (Nunc, cat. no. 140675) in 200 µL/cm^2^ Essential 6 (DMEM/F12 (ThermoFisher, cat. no. 11330-057), 64 mg/L ascorbic acid 2-phosphate (Sigma-Aldrich, cat. no. A92902), 14 μg/L sodium selenite (Sigma-Aldrich, cat. no. 214485), 543 mg/L sodium bicarbonate (Sigma Aldrich, cat. No. 55761), 20 μg/L Insulin (Sigma-Aldrich, cat. no. I9278), 10.7 μg/mL human recombinant transferrin (Sigma-Aldrich, cat. no. T3705)) with 10 µM Y-27632 (Tocris, cat. no. 1254) and cultivated at 37°C and 5% CO_2_ for 24 h. Between day 1-15, every 48 h media was changed to E6 with 200 µL/cm^2^ for maturation of iSMCs. On day 16, cells were detached with collagenase for 5 min at 37°C and cell-collagenase-mix was transferred into a tube and shaked at 70 rpm at 37°C for 5 min. Cell suspension was filtered with 100 µm pore size filters (Greiner, cat. no. 542100) and centrifuged at 485 g. Gel matrix was prepared on-ice in the following order for one 48-well (Nunc, cat. no. 150687): 110 µL collagen 13.75 µL PBS pH7.4 (10X) (Gibco, cat. no. 70011-036) incl. phenol red,16.25 µL 0.4 M NaOH, 137.5 µL cell suspension including 450.000 cells. A volume of 250 µL from the gel-cell-mix was added to one 48-well and solidified for 30 min at 37°C and 5% CO_2_. After solidification, gel-cell-disc was covered with 0.5 ml of warm E6 and cultivated at 37°C and 5% CO_2_ for 24 h. On day 17, media was changed to 0.5 ml E6 with additives: Activin A (Peprotech, cat. no. GMP120-14E, 0.25 ng/ml), sotatercept (MedChemExpress, HY-P99590, 10 µg/ml). The gel disc was released from the edges of the well with a 10 µl pipet tip. Cultivation was prolonged for another 72 h and shrinkage of gel disc was imaged via brightfield of the Axioscope 7 microscope (ZEISS).

### Bulk RNA-sequencing

Transcriptomics was performed by bulk RNA-sequencing. On day 0, fresh differentiated iSMCs were seeded as passage 1 at 15.000 cells/cm^2^ on vitronectin-coated (Gibco, cat. no. A31804) plates (Nunc, cat. no. 140675) in 200 µL/cm^2^ Essential 6 (DMEM/F12 (ThermoFisher, cat. no. 11330-057), 64 mg/L ascorbic acid 2-phosphate (Sigma-Aldrich, cat. no. A92902), 14 μg/L sodium selenite (Sigma-Aldrich, cat. no. 214485), 543 mg/L sodium bicarbonate (Sigma Aldrich, cat. No. 55761), 20 μg/L Insulin (Sigma-Aldrich, cat. no. I9278), 10.7 μg/mL human recombinant transferrin (Sigma-Aldrich, cat. no. T3705)) with 10 µM Y-27632 (Tocris, cat. no. 1254) and cultivated at 37°C and 5% CO_2_ for 24 h. On day 1 and 3, media was changed to E6 with 200 µL/cm^2^. On day 5, media was changed to 200 µL/cm^2^ E6 with additives: Activin A (Peprotech, cat. no. GMP120-14E, 0.25 ng/ml), sotatercept (MedChemExpress, HY-P99590, 10 µg/ml). On day 7, after 48 h, cells were detached with Accutase^TM^ (Gibco, cat. no. A1110501), lyzed with TRIzol Reagent (Invitrogen, cat. no. 15596018) and isolated with the Nucleospin RNA II Kit (Macherey-Nagel, cat. no. 740955) and stored at -80°C. Bulk RNA-sequencing was performed by Novogene, Germany.

### Principle component analysis

Principle component analysis (PCA) analysis for untreated samples was performed based on the Log2(TPM+1)-transformed expression counts using the PCA tools R package. The 2000 most variable genes were identified based on variance across samples and used for dimensionality reduction. Gene expression values were subsequently scaled per gene (row-wise z-score normalization) before PCA.

### SMC score analysis

To compare smooth muscle cell (SMC) marker expression scores in Vehicle-treated iSMCs with primary smooth muscle cells and undifferentiated hiPSCs, publicly available RNAseq data from human pulmonary artery smooth muscle cells (PASMCs; GSE144274) and human iPSC lines from the HipSci resource (HPSI0114, https://www.hipsci.org) were used, respectively. Lung vascular SMC marker genes were defined based on the gene set reported by Travaglini et al. (Table 1). For each sample, an SMC score was calculated as the mean expression of Travaglini SMC genes. For primary PASMC and hiPSC derived SMC samples, variance-stabilized expression values obtained using DESeq2 were used, and SMC scores were computed as the mean VST expression across SMC genes per sample. For external hiPSC samples, transcript-level abundance estimates in TPM were aggregated to gene-level expression based on HGNC gene symbols, and SMC scores were calculated as the mean log2(TPM + 1) expression across SMC genes per sample. SMC scores were compared across sample groups using violin and box plots to visualize the distribution of SMC gene-set expression.

**Table 1:**
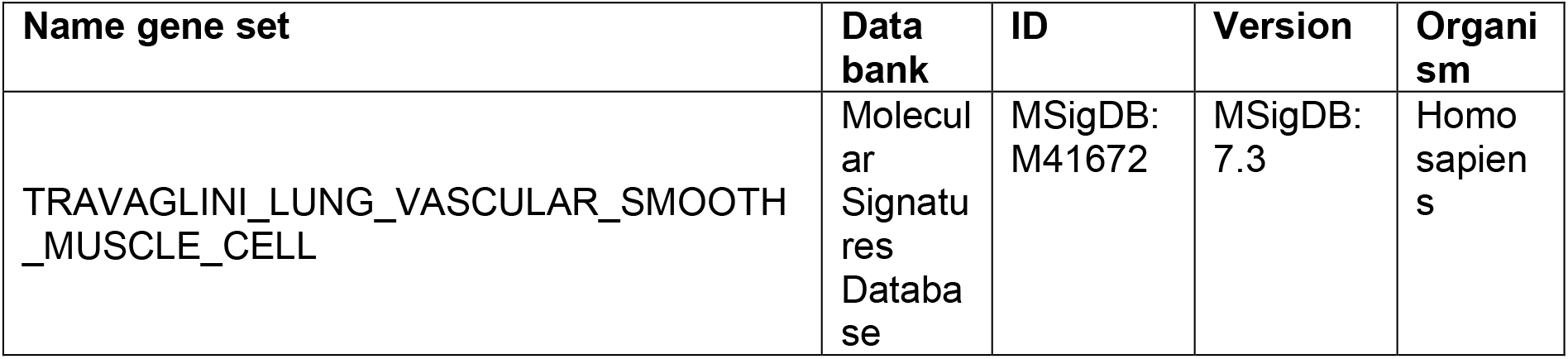
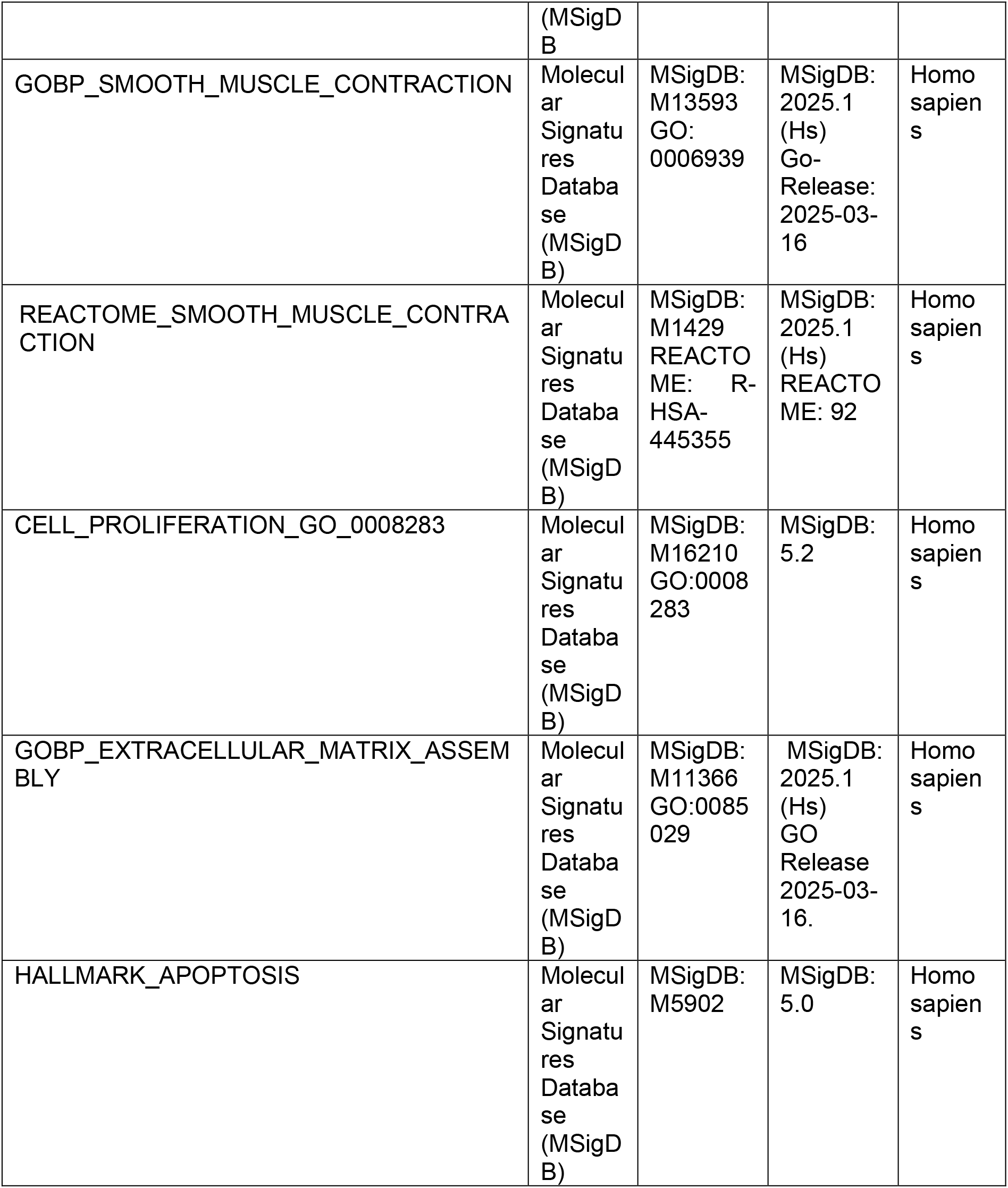
All listed gene sets are publicly available from the Molecular Signatures Database (MSigDB).

### Hallmark/marker gene set scoring

For hallmark gene set scoring, bulk RNAseq expression values were analyzed in R (v4.5.0) using a gene-by-sample FPKM matrix. Gene symbols were used as identifiers; when multiple rows mapped to the same gene symbol, values were merged by taking the mean FPKM per sample. Sample names were parsed into genotype, replicate, and treatment metadata to generate a sample annotation table. To visualize gene set-associated expression changes across conditions, samples were grouped by genotype and treatment, and gene expression was averaged across biological replicates within each group (row-wise mean across samples). For each genotype, treatment-induced gene expression level changes were quantified relative to the corresponding vehicle control by computing log2 fold-changes with a small pseudo-count (1 × 10⁻⁶) to avoid division by zero. For heatmap gene ordering, genes were classified into two predefined, manually curated subsets representing genes selected to be positively supporting or opposing the corresponding gene set or hallmark signature, where applicable. Within each predefined subset, genes were hierarchically clustered based on the fold-change matrix and the final gene order for heatmap visualization was obtained by concatenating the clustered order of the positively supporting gene subset followed by the opposing gene subset. Row-wise z-score normalization was performed across all displayed conditions or across each genotypes before visualization, genes with zero variance were assigned a z-score of 0. Heatmaps were rendered using ggplot2 geom_tile with a diverging colour scale centred at zero, and gene labels were annotated to indicate membership in the predefined gene subsets.

### Functional enrichment analysis

Functional enrichment analysis of the gene sets was performed using Enrichr. Cell type enrichment was evaluated using the CellMarker 2024. Enrichment results were visualized using bar plots generated by Enrichr ^97^.

### Myofibroblast signature scoring and expression comparison

A myofibroblast signature score “Myoscore” was calculated from the bulk RNA-seq FPKM expression matrix using a predefined, manually curated set of myofibroblast-associated marker genes with R (v4.5.0). Gene symbols were used as identifiers; if multiple rows mapped to the same gene symbol, values were merged by taking the mean FPKM per sample. After removing duplicated gene symbols from the marker list, only genes present in the expression matrix were retained. Myoscore distributions across groups were visualized using boxplots containing median, the interquartile range (IQR), and whiskers = 1.5 × IQR with overlaid jittered points representing individual biological replicates. For heatmap visualization, gene expression was averaged across biological replicates within each genotype–treatment group, and treatment-associated changes were quantified as log2 fold-changes relative to the corresponding vehicle control within each genotype, using a small pseudo-count (1 × 10⁻⁶) to avoid division by zero. Genes were hierarchically clustered based on the fold-change matrix to define the heatmap row order. Row-wise z-score normalization (z = (x − mean)/sd) was performed across all displayed conditions prior to visualization, genes with zero variance were assigned z = 0, and heatmaps were rendered using ggplot2 geom_tile with a diverging color scale centered at zero.

To validate Myoscore result, we generated a myofibroblast gene set that is less dependent on subjective manual selection. We leveraged three independently annotated human lung single-cell RNA-seq datasets: GSE135893, GSE136831 and the Munich cohort dataset from Mayr et al, 2021. In each dataset, author-provided cell type annotations were used to define cell identities. Within each dataset, differential expression analysis was performed comparing myofibroblasts against all other cell types using Seurat’s FindMarkers function. To ensure robustness to feature selection, marker detection was conducted both across all genes and restricted to highly variable genes, and the resulting myofibroblast-enriched genes (padj < 0.05 and avg_log2FC > 0) were merged and de-duplicated. Candidate genes were further filtered for expression prevalence, retaining only those detected in at least 30% of myofibroblast cells.

To quantify cell-type specificity, a specificity score was calculated for each candidate gene as the difference between its mean expression in myofibroblasts and the maximum mean expression observed across all non-myofibroblast cell types. This specificity metric was combined with the differential expression effect size to derive a composite ranking score (0.7 × specificity score + 0.3 × avg_log2FC), and the top 100 genes per dataset were selected as data-driven myofibroblast marker sets. Marker gene lists derived independently from the three datasets were then compared, and the final myofibroblast gene set was defined as genes identified in at least two of the three datasets, representing the most consistent cross-dataset signature. The myofibroblast gene set was subsequently used for module score-based validation in all three single-cell datasets to confirm its robust and selective enrichment in myofibroblast populations across datasets and analytical frameworks, after which this gene set was used for myofibroblast signature scoring as validation to the results previously generated.

## Data availability

All data will be made publicly available upon publication.

## Author contributions

A.S. performed and analyzed experiments and wrote the manuscript, L.C., L M., G.B., C.P., T.K., J.B. performed experiments, Y. W., C. V., A.K. performed data analysis, A. W. performed data analysis and provided scientific input, R.S., A.R. provided scientific input, J. C. K., M. M. H. provided patient samples and scientific input, RO, and UM conceptualized and supervised the study, provided scientific input and wrote the manuscript. All authors reviewed and approved the final version of the manuscript.

## Acknowledgments

The authors would like to thank G. Hansmann for his help with sample collection and N. Cleve for technical support.

