## Supplementary figures and images for "iPSC modeling of pulmonary arterial hypertension to uncover pathomechanisms and unrecognized modes of action of sotatercept"

### Folie6.PNG

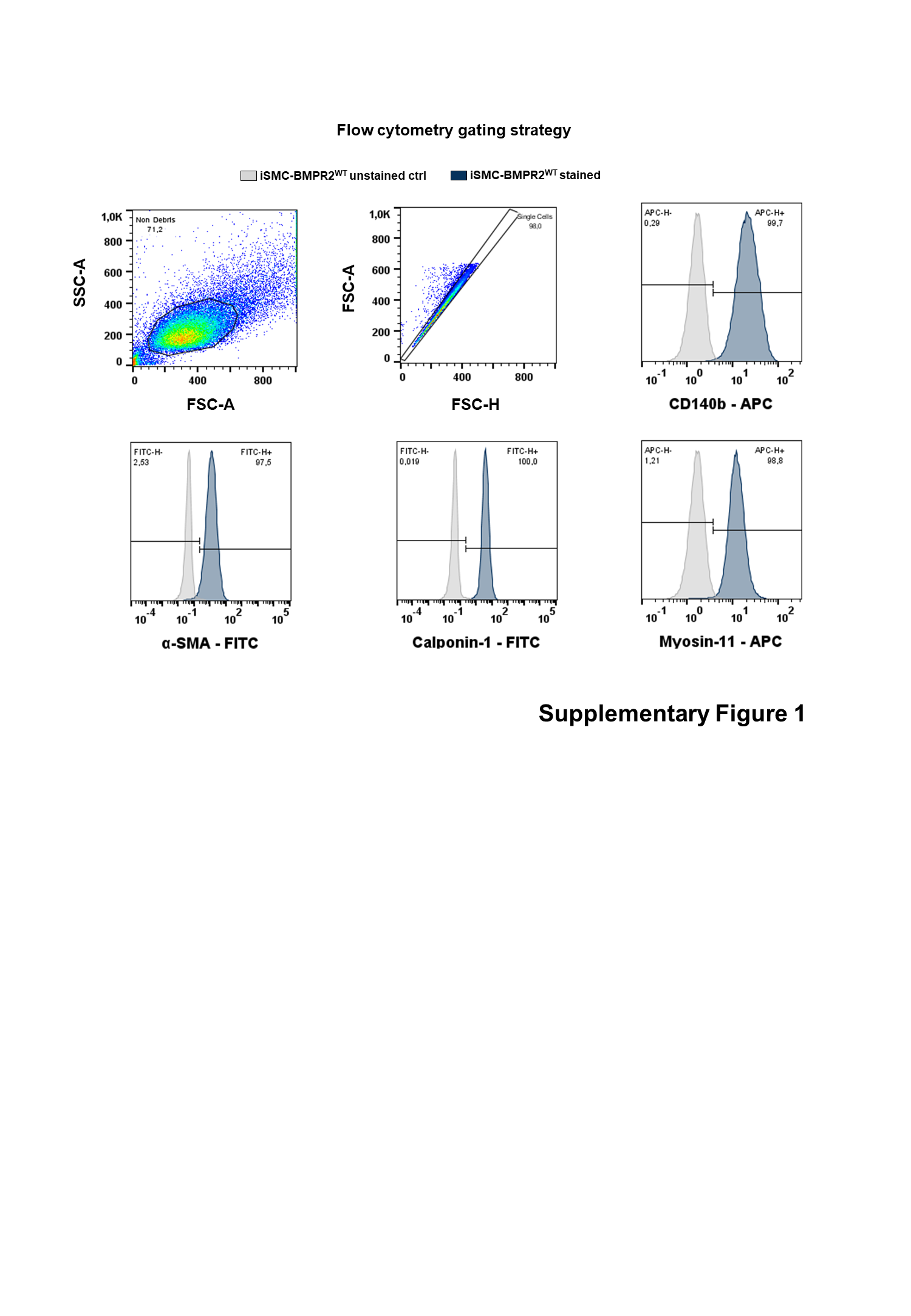

### Folie7.PNG

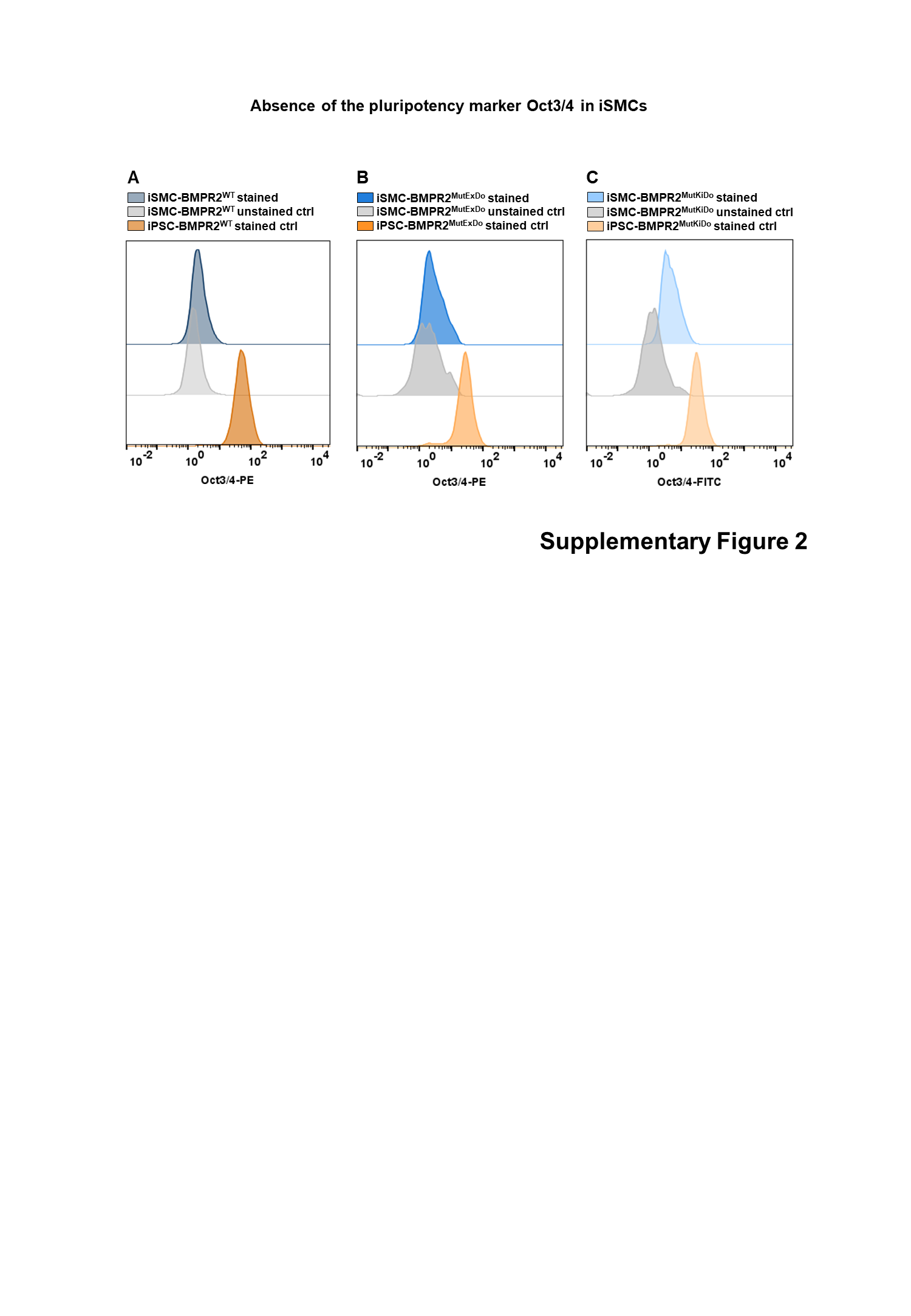

### Folie8.PNG

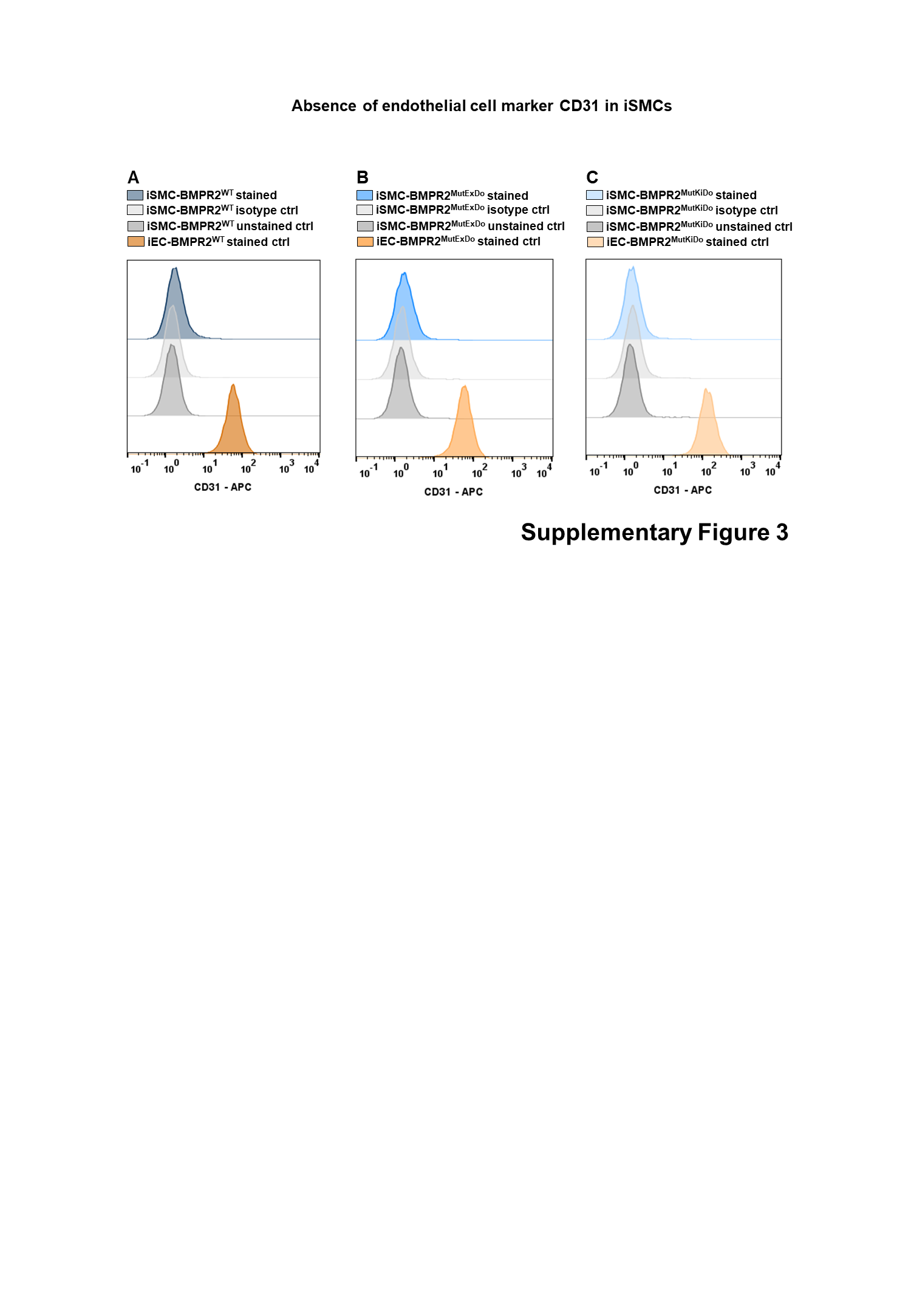

### Folie9.PNG

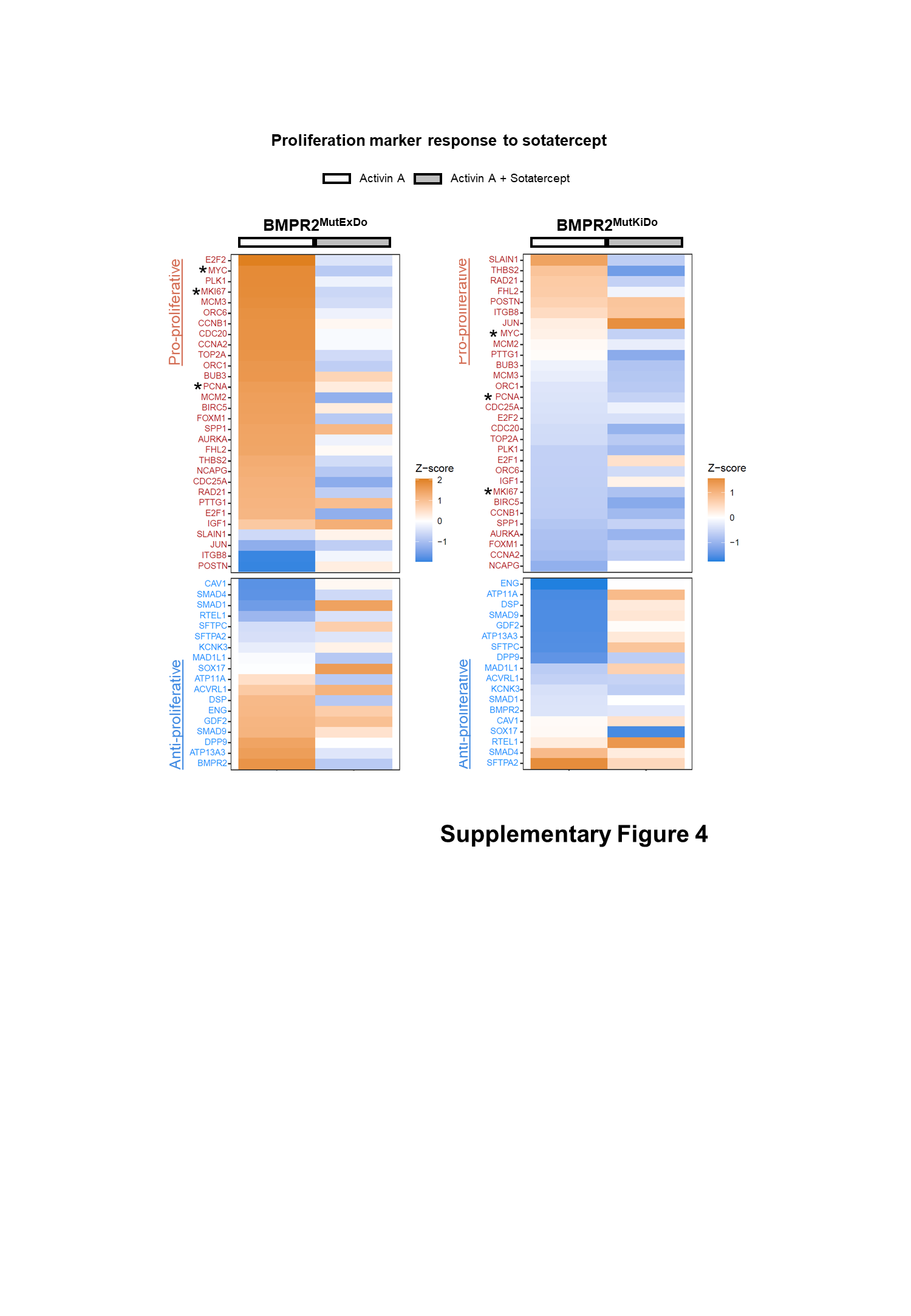

### Folie10.PNG

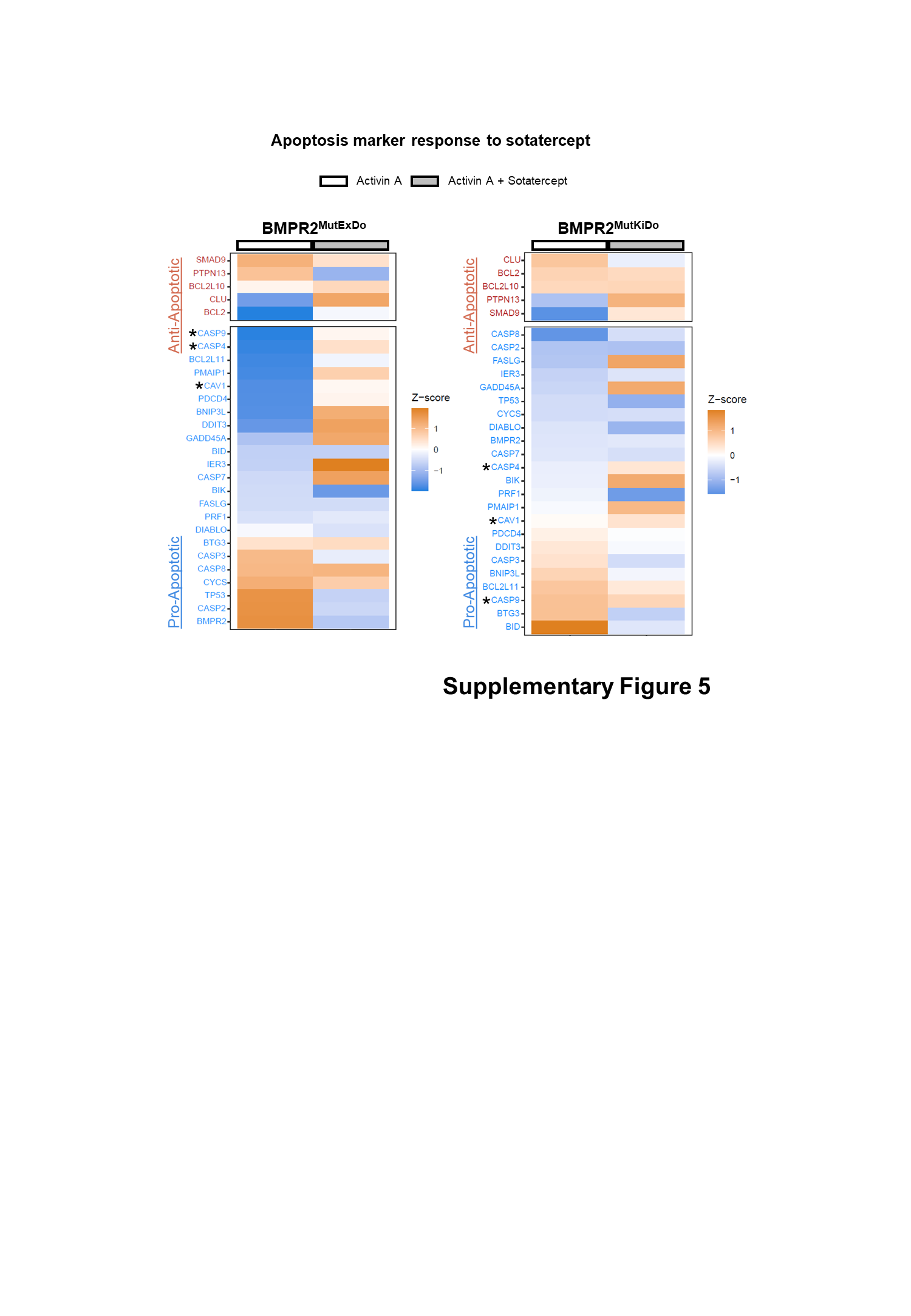

### Folie11.PNG

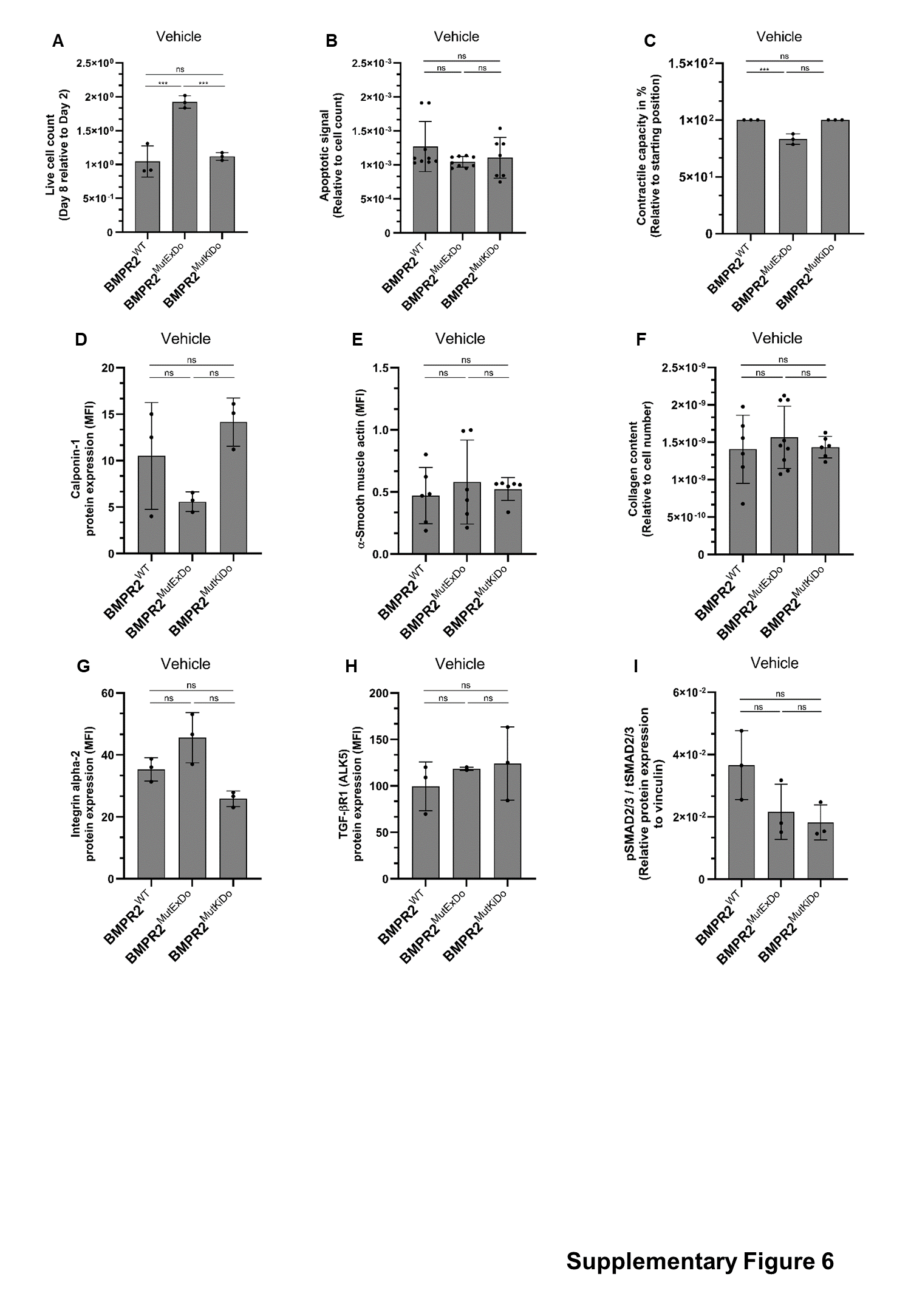

### Folie12.PNG

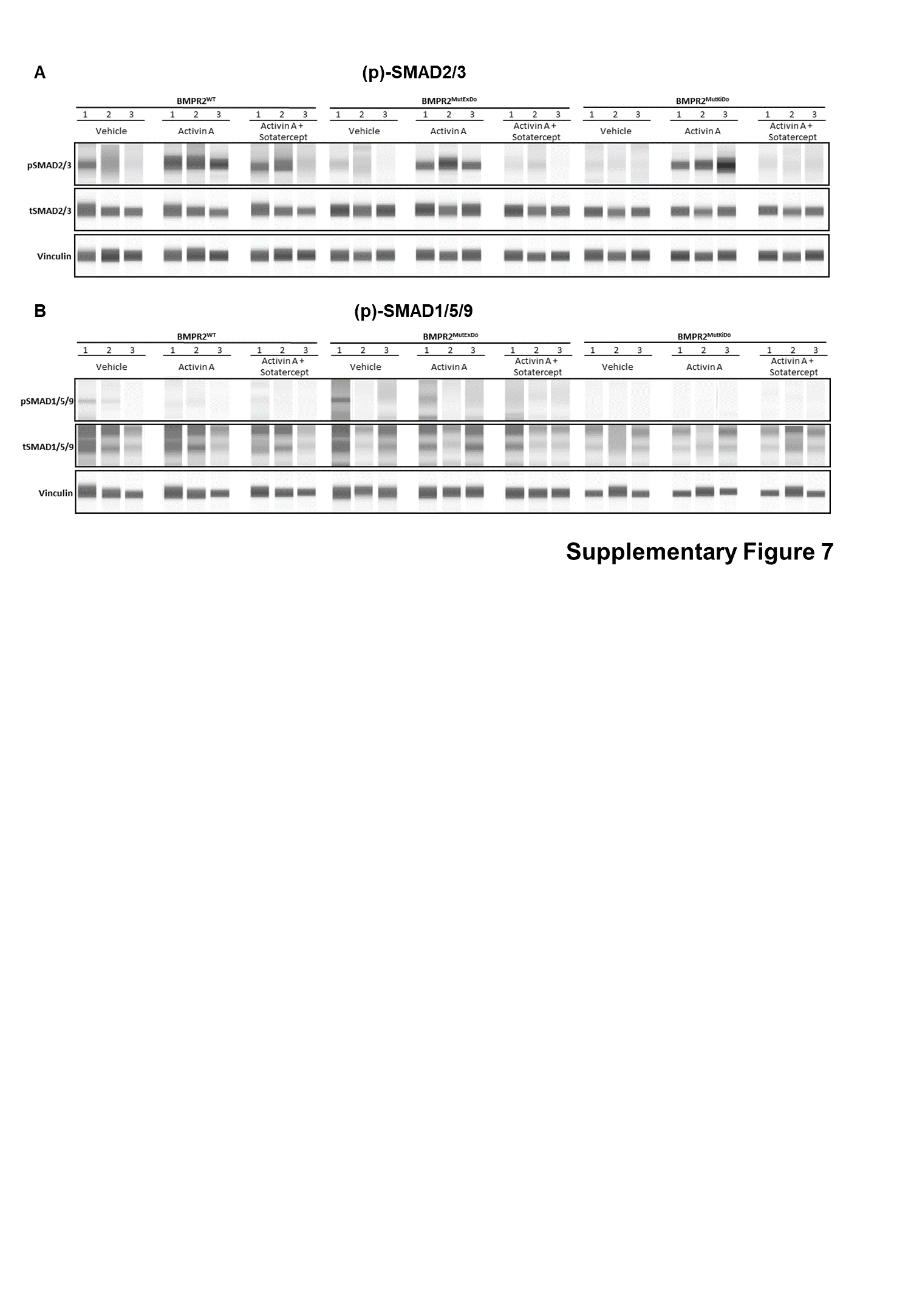

### Folie13.PNG

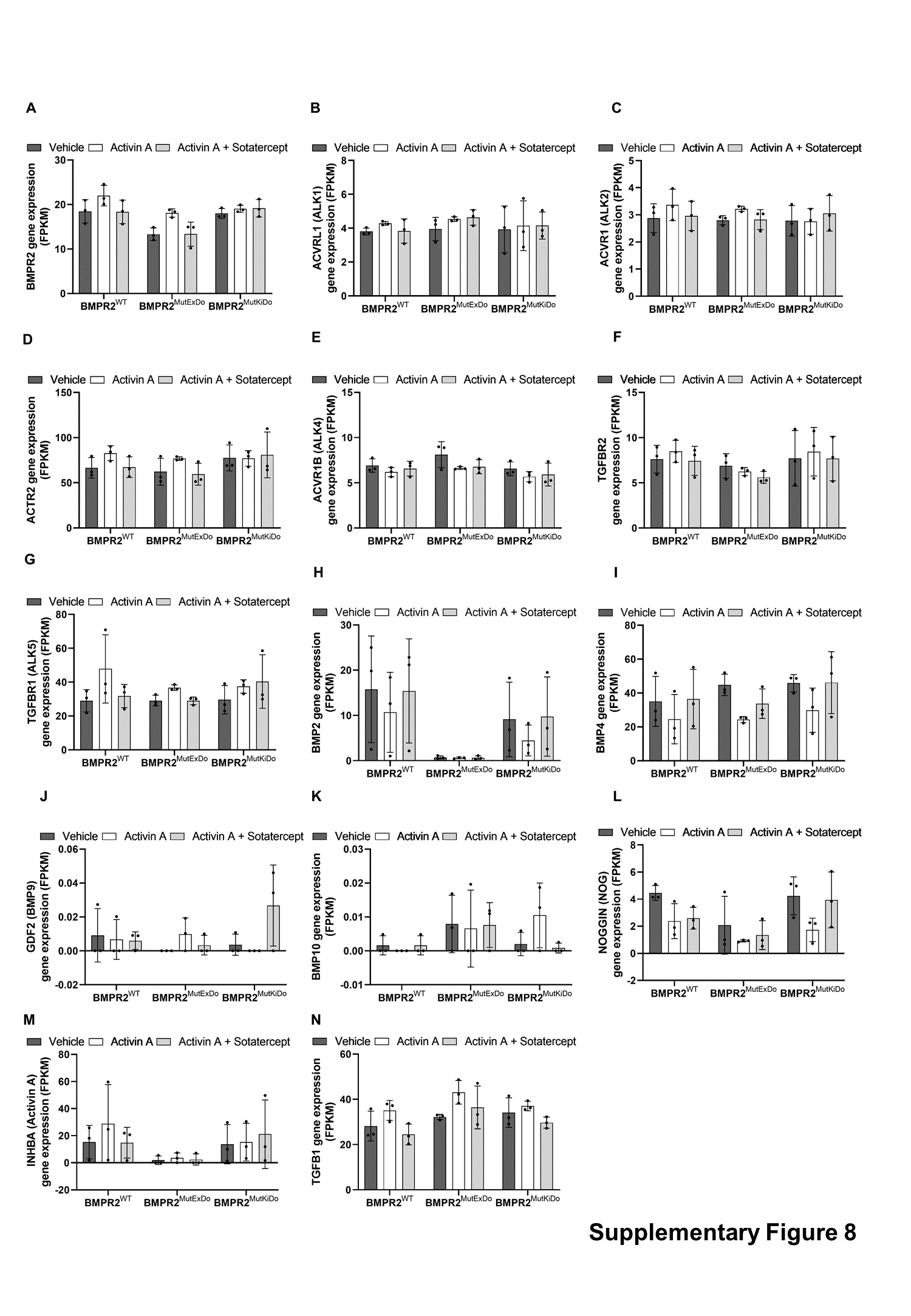
